# The *Mycobacterium tuberculosis* DNA-repair helicase UvrD1 is activated by redox-dependent dimerization via a 2B domain cysteine conserved in other Actinobacteria

**DOI:** 10.1101/2021.10.26.465901

**Authors:** Ankita Chadda, Drake Jensen, Eric J. Tomko, Ana Ruiz Manzano, Binh Nguyen, Timothy M. Lohman, Eric A. Galburt

## Abstract

*Mycobacterium tuberculosis (Mtb)* causes Tuberculosis and, during infection, is exposed to reactive oxygen species (ROS) and reactive nitrogen intermediates (RNI) from the host immune response that can cause DNA damage. UvrD-like proteins are involved in DNA repair and replication and belong to the SF1 family of DNA helicases that use ATP hydrolysis to catalyze DNA unwinding. In *Mtb,* there are two UvrD-like enzymes where UvrD1 is most closely related to other family members. Previous studies have suggested that UvrD1 is exclusively monomeric, however it is well-known that *E. coli* UvrD and other UvrD-family members exhibit monomer-dimer equilibria and unwind as dimers in the absence of accessory factors. Here, we reconcile these incongruent studies by showing that *Mtb* UvrD1 exists in monomer, dimer, and tetramer oligomeric forms where dimerization is regulated by redox potential. We identify a 2B domain cysteine, conserved in many Actinobacteria, that underlies this effect. We also show that UvrD1 DNA unwinding activity correlates specifically with the dimer population and is thus titrated directly via increasing positive (i.e. oxidative) redox potential. Consistent with the regulatory role of the 2B domain and the dimerization-based activation of DNA unwinding in UvrD-family helicases, these results suggest that UvrD1 is activated under oxidizing conditions when it may be needed to respond to DNA damage during infection.

## Introduction

DNA repair plays an essential role in the ability of organisms to maintain genome integrity in the face of environmental stresses. One particularly flexible and conserved pathway is Nucleotide Excision Repair (NER) which detects, and repairs bulky nucleotide lesions caused by UV light, environmental mutagens, and a subset of oxidative lesions (1–3). In bacteria, global genome NER is initiated when lesions are recognized directly by UvrA, although an alternative pathway called transcription-coupled NER depends on RNA polymerase stalling as the initiation event (4–7). The removal of the lesion eventually requires the recruitment of a helicase to the site of damage. In Eukaryotes, this function is filled by TFIIH (8–10), while prokaryotes utilize the UvrD-family enzymes (1,3, 11). In addition to its role in NER, UvrD participates in a range of other pathways of DNA metabolism such as replication (12–15) and recombination (16–18).

UvrD has been well-characterized in many contexts and from model organisms including *E. coli* and *B. subtilis.* It is a superfamily 1A (SF1A) helicase, as defined by core helicase domains 1A and 1B coupled with auxiliary 2A and 2B sub-domains (19, 20). It can both translocate on singlestranded DNA (ssDNA) and unwind double-stranded DNA (dsDNA) under specific conditions. More precisely, while monomers of UvrD-family members (UvrD, Rep and PcrA) are ATP-dependent ssDNA translocases, dimeric forms of these enzymes are required to unwind duplex DNA *in vitro* in the absence of accessory factors or force (21–27). In Rep, this activation is regulated by the mobile 2B domain as both deletion of the 2B domain or a crosslinked 2B domain construct activate the Rep monomer for unwinding (28, 29). Activation of the dimeric UvrD helicase is also accompanied by re-orientation of its 2B sub-domain (30). Additionally, the rotational orientation of the 2B domain regulates the force-dependent unwinding activity of both UvrD and Rep monomers (31, 32). Helicase activation can also occur via binding with accessory factors. For example, *B. stearothermophilus* RepD activates PcrA monomers (31, 32, 25) and the mismatch repair protein MutL activates UvrD monomers (33, 34). Furthermore, these interactions directly affect the orientation of the regulatory 2B domain (34). In addition to its association with other repair proteins (32), UvrD associates with RNA polymerase through its C-terminal RNAP Interaction Doman (RID) during one mode of transcription coupled NER (35–37). This interaction leads to the stimulation of RNAP backtracking and the recruitment of UvrAB (37).

While many studies have been reported focusing on UvrD-family helicases from model bacteria, less is known about these enzymes in the distantly related human pathogen, *Mycobacterium tuberculosis (Mtb). Mtb* is the causative agent of Tuberculosis and is the leading cause of death worldwide from an infectious agent (38). Although DNA metabolism pathways such as transcription and repair are generally conserved in bacteria, important differences exist (39–42). This appears to be especially true in *Mtb,* perhaps as it is highly evolved for a relatively narrow niche (43). Interestingly, and in contrast to model bacteria, *Mtb* contains two UvrD family enzymes: UvrD1 and UvrD2 (44, 45). UvrD1 has high homology to *E. coli* UvrD including the C-terminal RID (45, 46). Previous work on *Mtb* UvrD1 has shown that it is important for survival after UV and oxidative damage as well as for pathogenesis in mice (47). In stark contrast to other UvrD-family members, UvrD1 has been reported to be monomeric and to either possess helicase activity directly or require activation via the binding of *Mtb* Ku (45, 46, 48).

Here we report that UvrD1 exists in monomer, dimer, and tetrameric forms where dimerization is redox-dependent and is correlated with helicase activity. We identify a 2B domain cysteine that is required for the redox-dependent dimerization, demonstrating that the 2B subdomain is directly involved in dimerization. Our results explain the function of UvrD1 in the context of the large body of work on UvrD-family proteins and suggest a model where UvrD1 senses the oxidative conditions within human macrophages during infection through dimerization, resulting in activation of its DNA unwinding activity needed for DNA repair and other DNA metabolic pathways (49–51).

## Results

### The oligomeric state of UvrD1 is redox-dependent

Previous studies reported that UvrD1 exists exclusively as a monomer in solution (45, 48). However, upon purifying UvrD1 as described in the Methods, we observed two elution peaks from an S300 size exclusion column run at 4 °C in Tris pH 8.0 at 25 °C, 150 mM NaCl, 10% glycerol, and no DTT, consistent with the molecular weights of both monomer (85 kDa) and dimer (170 kDa) species **(Fig. S1)**. This result is consistent with studies of *E. coli* UvrD, which exhibits a monomer-dimer-tetramer equilibrium (23, 26). To examine this more quantitatively, we performed analytical ultracentrifugation sedimentation velocity experiments in TRIS pH 8.0 at 25 °C and 20% glycerol (from here on defined as Buffer A) with 75 mM NaCl and 2.5 μM UvrD1. The continuous sedimentation coefficient (c(s)) distribution (52) shows three peaks that we assign to monomer, dimer and higher order oligomers (**Fig. 1A, Supplemental Table 1)**. The positions of the peaks do not change with UvrD1 concentration indicating that each peak represents a single species; however, the amplitudes of the three peaks change with UvrD1 concentration as expected for a self-assembling monomer-dimer-oligomer system **(Fig. S2)**. The oligomeric states of *E. coli* UvrD depend on the salt and glycerol concentrations, with the monomer population favored by higher salt and glycerol concentrations (26). We examined the salt dependence of the UvrD1 oligomeric state by sedimentation velocity at 2.5 μM UvrD1 in a range of NaCl concentrations between 75 – 750 mM. Surprisingly, the ratio of monomer to dimer was relatively constant throughout the salt titration apart from the lowest salt concentrations where higher order oligomers were populated at the expense of the monomer population **(Fig. 1B, S3)**.

**Figure 1:**
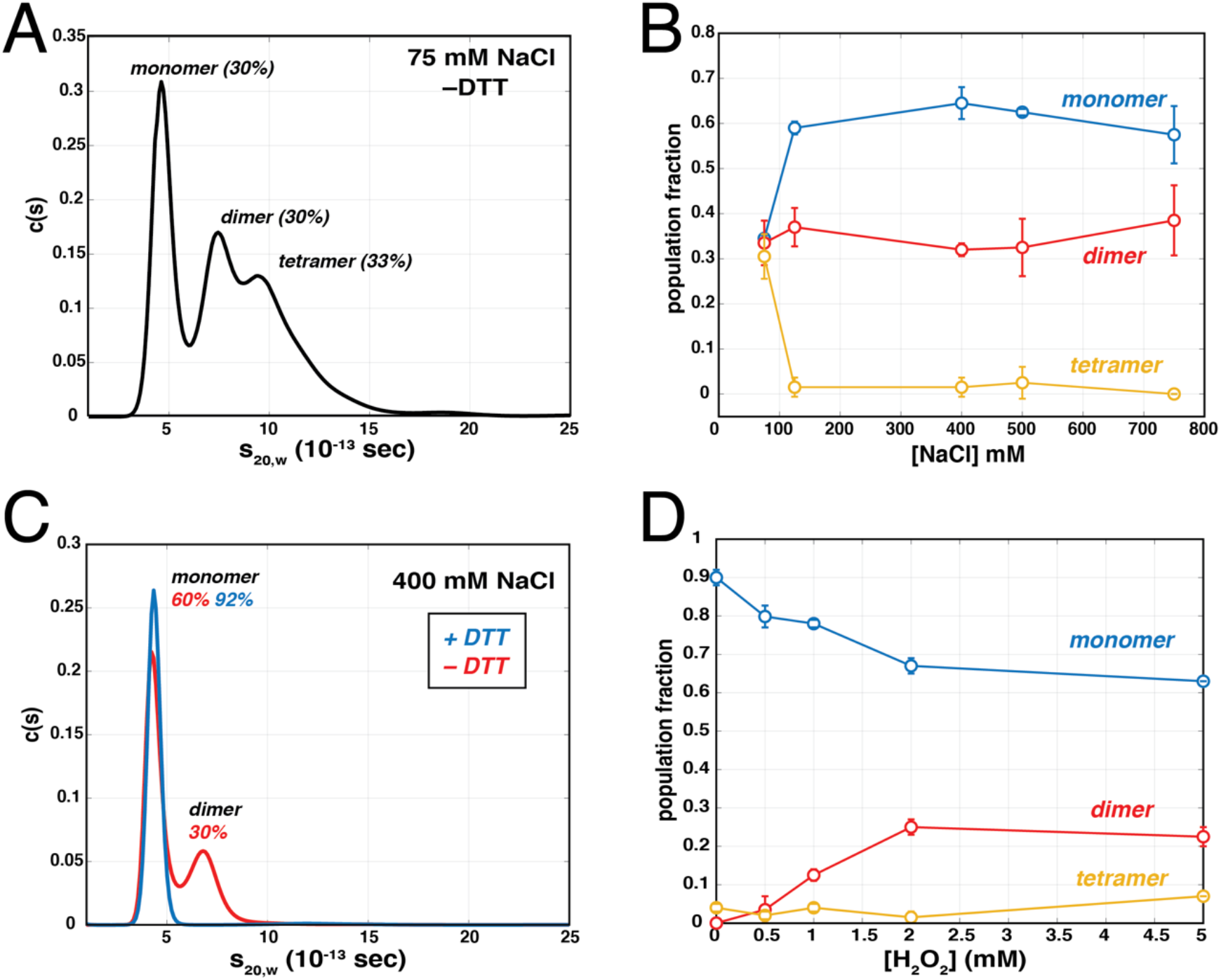
Oligomeric states of UvrD1. **(A)** Sedimentation velocity trace measured at 230 nm in Buffer A plus 75 mM NaCI in the absence of DTT reveals the presence of monomeric, dimeric, and tetrameric species. **(B)** Summary of results from AUC velocity experiments using 2.5 μM UvrD1 in the absence of DTT as a function of NaCl concentration **(Fig. S3)**. From 75 mM to 750 mM NaCl, the fraction of monomer present in the monomeric state (blue), dimeric state (red), and tetrameric states (yellow) are shown. **(C)** The continuous distribution of species from AUC velocity runs measured at 280 nm with 400 mM NaCl in the presence and absence of 1 mM DTT. **(D)** After treatment of 2.5 μM WT UvrD1 in 75 mM NaCl with 1 mM DTT, a titration series of H_2_O_2_ from 0 mM to 5 mM was run in AUC velocity experiments **(Fig. S5)**. Increasing concentrations of H_2_O_2_ result in a decrease in the fraction found in the monomeric state (blue) and an increasing fraction found in the dimeric state (red). Higher order oligomeric states (yellow) represented less than 10% under all conditions.

In contrast, the addition of 1 mM DTT at 400 mM NaCl in Buffer A shifted the species fraction dramatically to favor the monomer (96%, **Fig 1C**). Even at the lowest NaCl concentration of 75 mM NaCl, the addition of 1 mM DTT resulted in a nearly uniform monomer population (90%, **(Fig. S4)**. Thus, we hypothesized that UvrD1 dimerization is dependent on redox potential and reasoned that oxidative conditions should favor dimer formation just as reductive conditions favor the monomer species. To test this, we performed sedimentation velocity experiments in the presence of oxidizing agents after reduction by 1 mM DTT. As predicted, titration of hydrogen peroxide (H_2_O_2_) resulted in an increase in the dimer population **(Fig. 1D, S5)**. At 2 mM H_2_O_2_, the population fraction of dimer saturated at ~25% and the addition of more hydrogen peroxide did not lead to more dimer formation.

### A 2B domain-2B domain disulfide bond is responsible for redox-dependent dimerization of UvrD1

The dependence of oligomerization on oxidation suggested a role for a thiol-containing amino acid such as methionine or cysteine. In particular, we considered that the potential of cysteines to form disulfide bonds could lead to the formation of dimeric and higher order oligomers. UvrD1 has three cysteine residues for which we estimated their approximate position by generating a threaded homology model of UvrD1 based on the structure of *E. coli* UvrD (PDB:3LFU, PHYRE2, Methods, **Fig. 2A**). This model shows that, while two cysteines appear buried within the 1A and 1B domains (C107 and C269), a third cysteine (C451) is surface exposed within the 2B domain. Surface calculations of our model with Chimera (53) confirmed this as only C451 possesses solvent exposed surface area in both open (based on *E. coli* PDB:3LFU structure) and closed (based on *G. stearothermophilus* PDB:3PJR structure) conformations **(Fig. S6)**. We hypothesized that a disulfide bond between the 2B cysteines of two monomers was responsible for the redox-dependent dimerization. To test this, we constructed and purified a C451A mutant (which will we refer to as the 2B mutant) and examined whether it is able to form dimers. During purification of this construct, the S300 elution profile showed only a single peak consistent with a monomer in contrast to WT UvrD1 Sedimentation velocity experiments confirmed this result, as the 2B cysteine mutant was monomeric in both the presence and absence of DTT **(Fig. 2C, Supplemental Table 1).** In contrast, a double mutant of the 1A and 1B domain cysteines (C107T/C269T, now referred to as the 1A1B double mutant) maintained the ability to form dimers **(Fig. 2D, Supplemental Table 1)**.

**Figure 2:**
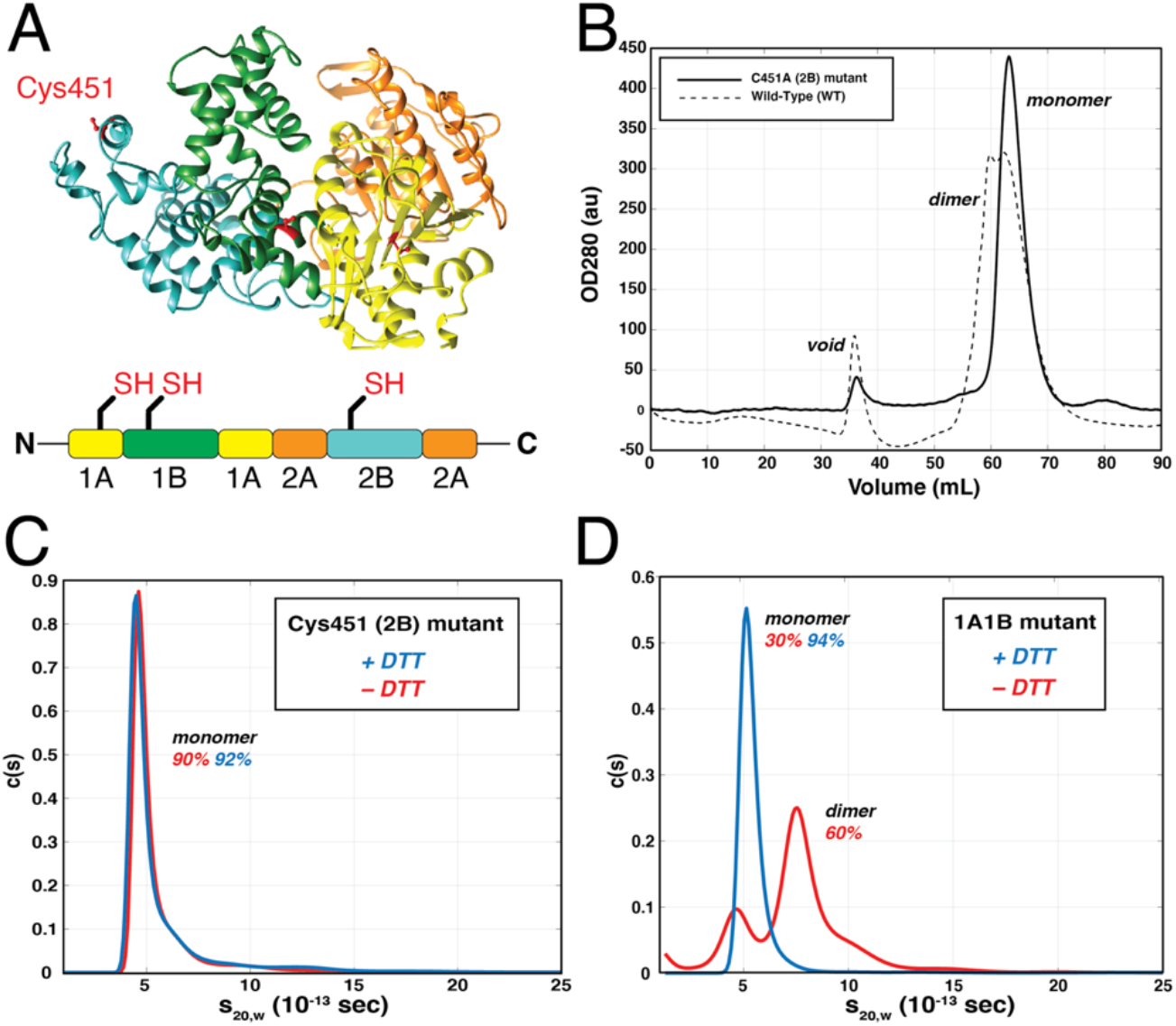
Redox-Dependence of UvrD1 dimerization is due to C451 in the 2B domain. **(A)** Predicted structure of UvrD1 from threading UvrD1 sequence on the E. coli UvrD structure (PDB: 3LFU). Domain organization is indicated as well as the position of the three cysteine residues described in the text. **(B)** Size exclusion chromatography (S300) of C451A mutant (solid) as compared to WT (dashed). Both constructs were run in Tris pH 8.0, 150 mM NaCl, 10% glycerol, and the absence of DTT as in Fig. S1. **(C)** AUC velocity experiments on the 2B mutant in Buffer A plus 75 mM NaCl and the presence (blue) and absence (red) of DTT indicate that the mutant loses the ability to dimerize. Each trace is an average of two runs and the population fractions for monomer and dimer are indicated. **(D)** In contrast, AUC velocity of the 1A1B double mutant in 75 mM NaCl and the presence (blue) and absence (red) of 1 mM DTT indicates that this mutant retains and even enhances dimer formation. Each trace is an average of two runs and population fractions for monomer and dimer are indicated.

The three cysteine residues found in UvrD1 are not conserved in *E. coli* UvrD, despite the presence of six cysteine residues **(Fig. 3A)**, and *E. coli* UvrD does not display a redox-dependent dimerization However, the 2B domain cysteine identified here is conserved across various Actinobacterial classes **(Fig. 3B,C, S8)**. A particularly high, but not universal, conservation was found in the Corynebacteriales order which includes *Mtb* and other pathogenic bacteria (54, 55). In addition, we found the same sequence in the PcrA helicase found in one strain of the Firmicute, *Clostridioides difficile* (NCTC13750), which represents another important human pathogen that interacts with macrophages (56, 57).

**Figure 3:**
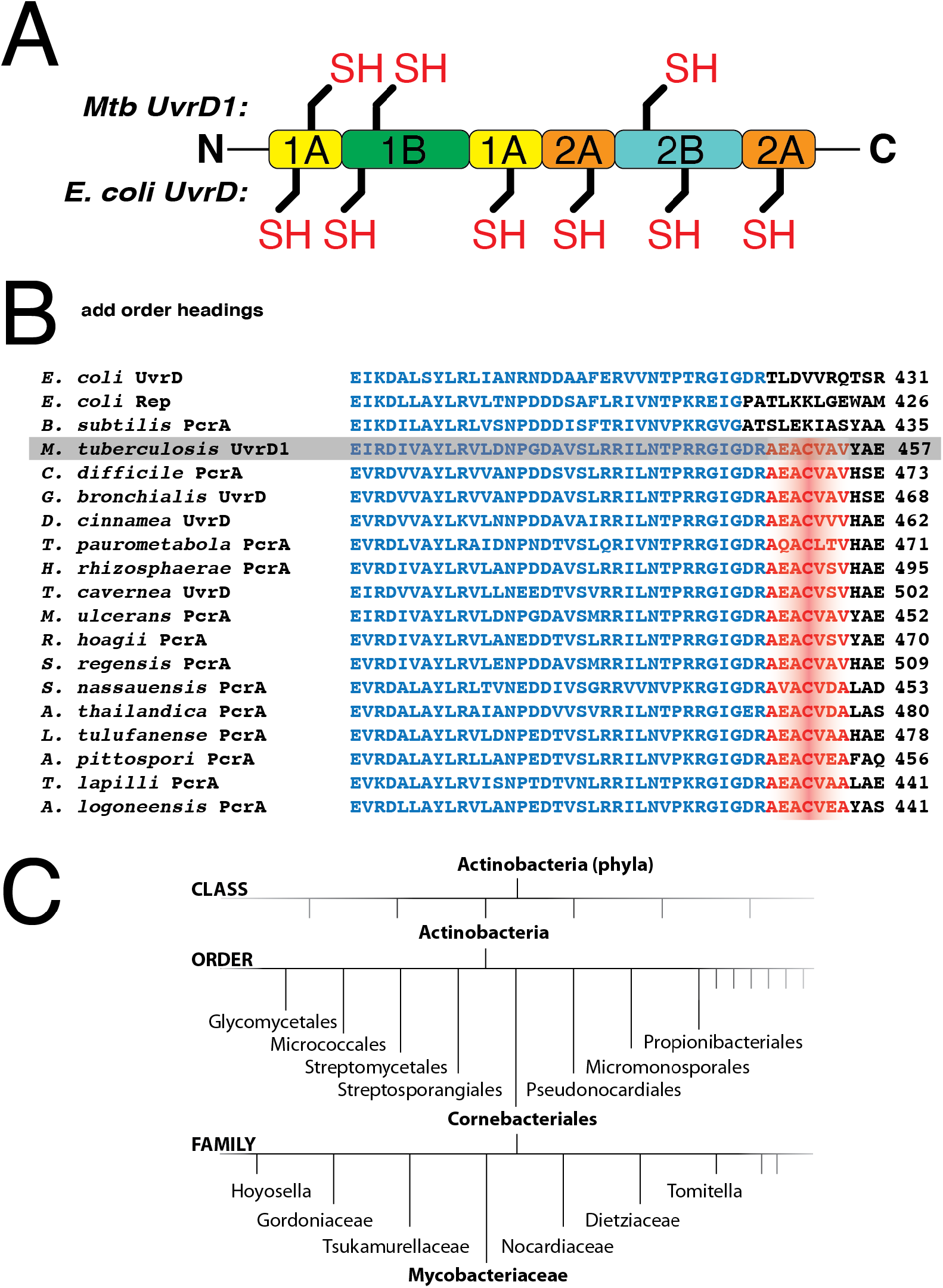
Sequence distribution across bacterial species. **(A)** Distribution of cysteine residues in *Mtb* UvrD1 compared to *E. coli* UvrD. **(B)** Sequence alignment of the 2B domain region containing C451. The blue sequence is conserved across all UvrD-like family members while the sequence containing 2B cysteine residue (red) is distinct from *E. coli* and B. subtilis UvrD family enzymes but conserved in many Actinomycetes. See Supplemental information for full species names. **(C)** Orders and Families of Actinobacteria where the 2B cysteine can be found in available sequence data.

### The dimer of UvrD1 is required for DNA unwinding activity

Previous studies of UvrD1 suggested that the monomer possesses helicase activity (48). However, this conclusion was based on measurements of helicase activity performed in solution conditions distinct from those used to examine its oligomerization state. In particular, the analysis of oligomeric state was performed in the presence of 5 mM DTT while helicase assays were performed in buffer lacking DTT entirely (48). In other studies, UvrD1 was surmised to be a monomer based on sedimentation through a glycerol gradient and was reported to have unwinding activity that was dramatically activated in the presence of *Mtb* Ku (45).

Given the observations of other UvrD-family helicases (23, 25, 26, 28, 58), we hypothesized that only the dimer of UvrD1 would be capable of unwinding DNA. To test this hypothesis, we used a stopped-flow assay to measure the time-dependence of UvrD1-catalyzed DNA unwinding **(Fig. 4A)**. Specifically, we used a double-stranded 18 bp DNA with a singlestranded dT_20_ 3’ flanking region (tail), with a Cy5 fluorophore on the 5’ end of the tailed strand, and a black hole quencher (BHQ2) on the 3’ end of the complementary strand as described previously (33, 34). In the double-stranded form, fluorescence from Cy5 is quenched due to the presence of the BHQ2 (59). Upon full unwinding, the strands are separated and the BHQ2 strand is trapped via an excess of unlabeled complementary “trap” DNA resulting in an increase in Cy5 fluorescence. This excess of trap also serves to bind any UvrD1 that dissociates from the labeled template ensuring single-round turnover conditions **(Fig. S9A)**. UvrD1 was pre-bound to the labeled DNA template and was loaded in one syringe. This solution was rapidly mixed with the contents of the other syringe consisting of an excess of trap strand, 5 mM Mg^+2^, and 1 mM ATP. The fraction of DNA unwound as a function of time was calculated by comparing experimental traces to a positive control consisting of fully single-stranded Cy5-labeled DNA and a negative control in the absence of ATP **(Fig. S9B)**.

**Figure 4:**
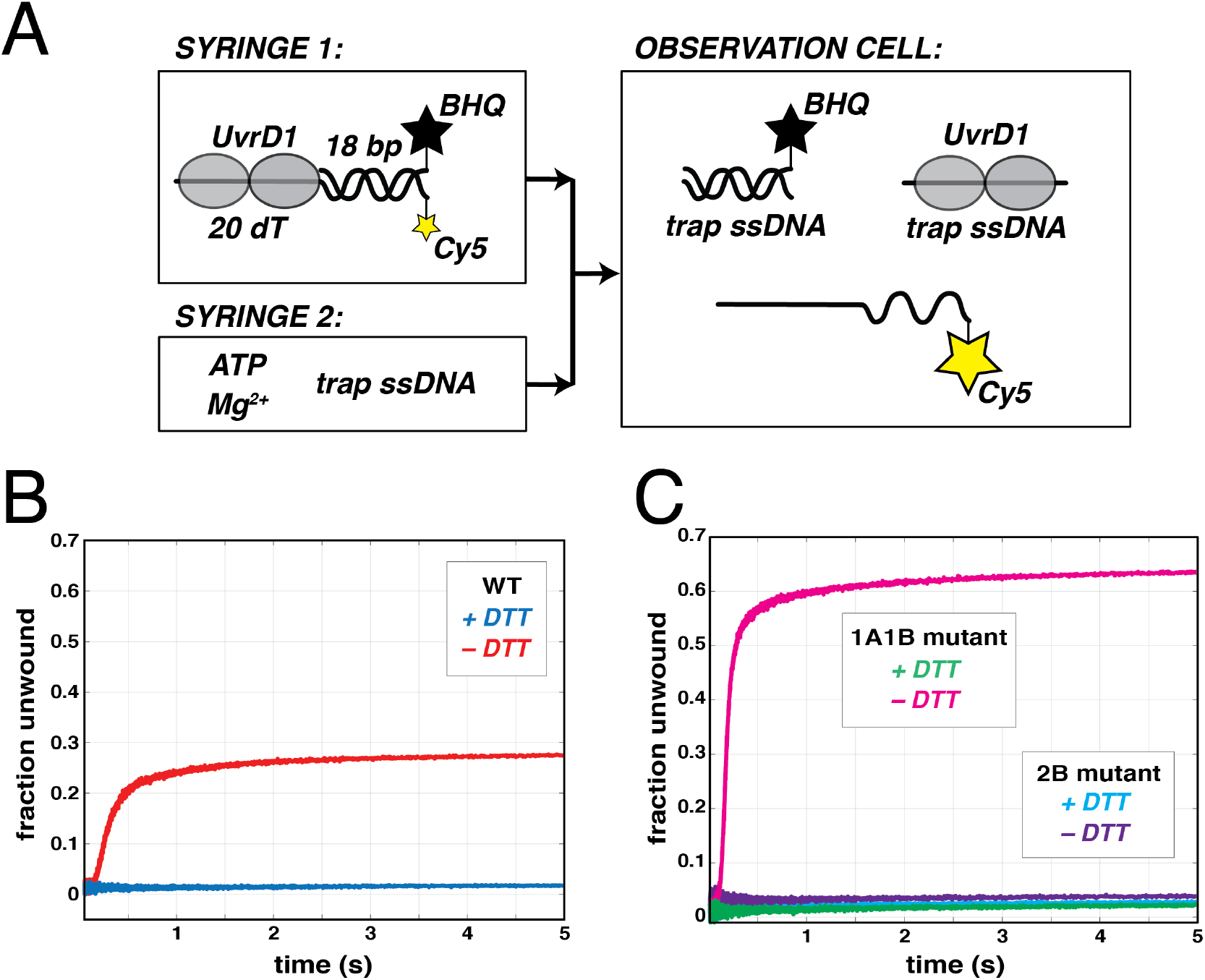
Unwinding activity is dependent on redox-dependent dimer. **(A)** Stopped-flow assay for monitoring DNA unwinding. One syringe is filled with UvrD1 that has been pre-equilibrated with a fluorescently labeled unwinding template consisting of an 18 bp duplex with a 3’ single-stranded 20 dT tail. The 5’ end of the loading strand is labeled with Cy5 and the 3’ end of the other strand is labeled with a black hole quencher (BHQ). A second syringe is filled with ATP, Mg^2+^, and single-stranded trap DNA. Upon mixing UvrD1 unwinds some fraction of the DNA resulting in a fluorescence enhancement due to the separation of the Cy5 from the BHQ. The excess trap DNA binds both the BHQ labeled single-strand and any free UvrD1 to establish single-round turnover conditions. **(B)** Fraction of DNA template unwound with 200 nM wild-type UvrD1 in the presence (blue) and absence (red) of DTT. **(C)** Fraction of DNA template unwound with 200 nM 1A1B double and 2B mutants of UvrD1 in the presence (green and cyan) and absence (pink and purple) of DTT in Buffer A with 75 mM NaCl.

In the absence of reducing agent, 200 nM WT UvrD1 can unwind ~27% of 2 nM duplex DNA in a single-round reaction **(Fig. 4B, red)** consistent with the fraction dimer in these conditions **(Fig. 1A)**. In addition, the kinetics and final percent unwound show a duplex-length-dependence as is expected **(Fig. S10A)**. Fits to an n-step model with a non-productive fraction **(Fig. S10A,B)** lead to an average unwinding rate of 64.8 ± 6.4 bp/s **(Supplemental Table 2)**. Similarly, a lag time analysis (60, 61) yielded an estimate of the unwinding rate of 83.3 ± 12.3 bp/s **(Fig. S10C,D)**. In contrast, in the presence of 1 mM DTT, which shifts the UvrD1 population to favor monomers, no unwinding is observed consistent with the UvrD1 dimer being required for unwinding activity **(Fig. 4B, blue)**. In addition, the 2B mutant (C451A), which we have shown is an obligate monomer even under oxidative conditions, lacks helicase activity in both the presence and absence of DTT **(Fig. 4C, cyan and purple)**. Consistently, the 1A1B double mutant is still able to unwind DNA in the absence of DTT **(Fig. 4C, pink)**. In fact, the 1A1B double mutant unwinds a higher fraction of DNA compared to WT (~63%), consistent with the AUC results suggesting that a higher fraction of the enzyme is in the dimeric form in the absence of DTT **(Fig. 2D)**. Thus, the dimer fraction of UvrD1 formed via the 2B domain disulfide bond is required for DNA unwinding activity.

### Both monomers and dimers of UvrD1 bind DNA

When considering why UvrD1 dimerization is required for DNA unwinding, we initially considered two hypotheses. One was that monomers are unable to bind DNA at the concentrations utilized and the second was that dimerization is required for helicase activation. To determine the nature of the DNA-bound species, we used sedimentation velocity to examine both WT UvrD1 and the 2B mutant in the absence of reducing agent and in the presence of fluorescently labeled 18 bp dT_20_ DNA. The results show that both monomers and dimers interact with the DNA and appear to bind the DNA in the same ratio as the free oligomeric forms. Specifically, in the presence of 1.5 μM WT UvrD1 and 1.5 μM DNA, 30% of the DNA was bound by monomers and 31% was bound by dimers **(Fig. 5A, red, Supplemental Table 3)**. In the presence of 1.5 μM of the 2B mutant, only monomers were bound to DNA **(Fig. 5A, blue)**. Even at higher molar ratios of monomeric UvrD1 (i.e., the 2B mutant) to DNA, only monomers are observed bound to DNA suggesting that DNA binding alone does not stimulate dimerization **(Fig. S11)**. In contrast, when WT UvrD1 in the absence of DTT is mixed with DNA possessing a shorter single stranded extension (dT_10_), only monomers bind **(Fig. 5B, Supplemental Table 3)** and no DNA unwinding is observed **(Fig. S12)**. This result is consistent with a model where each individual monomer interacts with the ssDNA in the context of the bound dimer as is seen with *E. coli* UvrD (23). Fluorescent anisotropy experiments with the unwinding substrate DNA (18 bp dT_20_) and either WT or 2B mutant UvrD1 yielded similar concentration dependencies of binding, consistent with the idea that the affinities of the monomeric and dimeric species are similar **(Fig. 5C)**.

**Figure 5:**
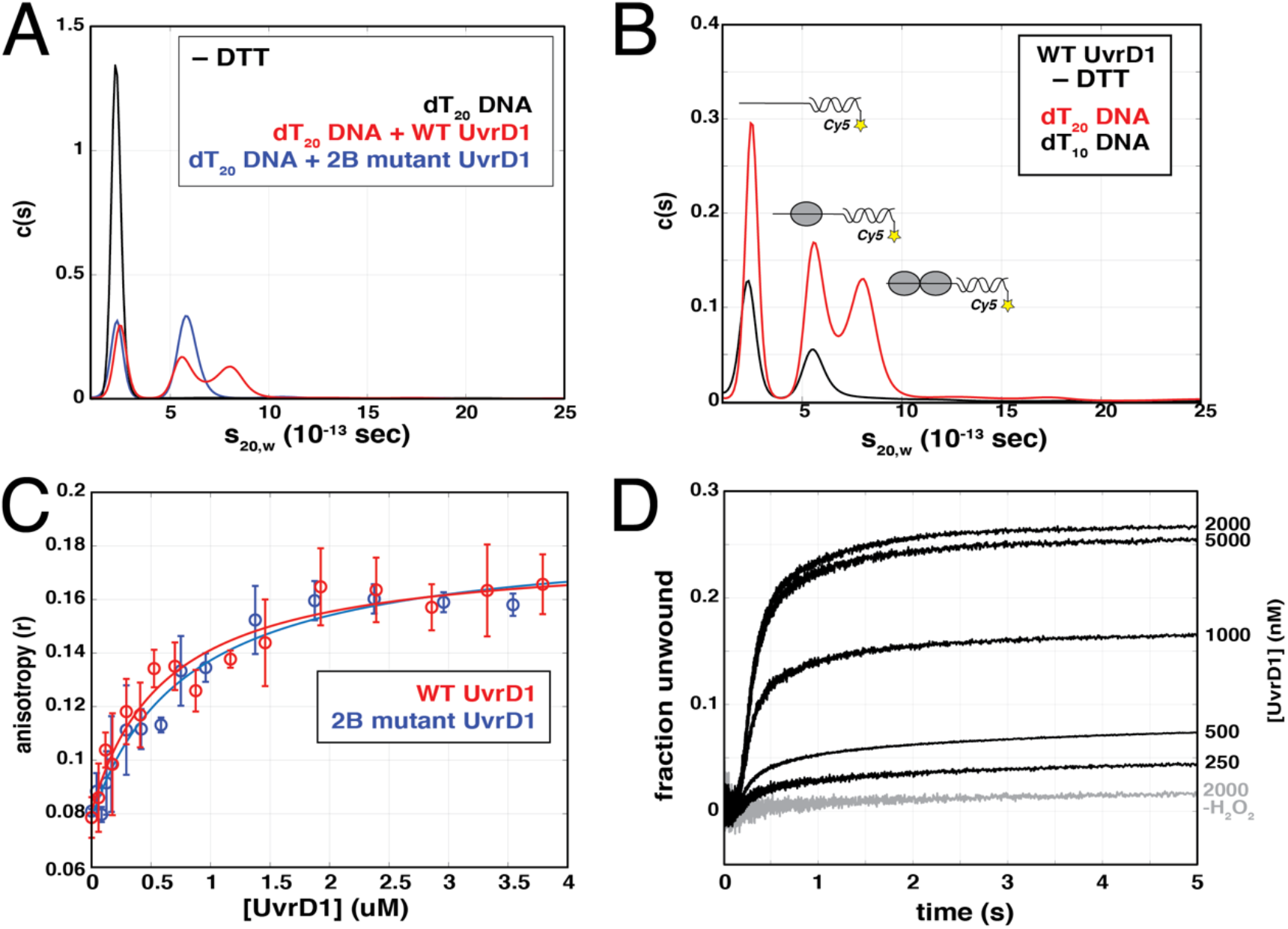
Monomer and dimer forms of UvrD1 both bind DNA unwinding template. **(A)** All experiments were performed in Buffer A with 75 mM NaCl. A Cy5-labeled DNA was used in sedimentation velocity experiments to specifically interrogate DNA-bound species. dT_20_ DNA alone (black), DNA and wild-type UvrD1 in the absence of reducing agent (red), and DNA and the 2B mutant in the absence of reducing agent (blue) are shown along with the bound species represented by each peak. **(B)** WT UvrD1 in the absence of reducing agent bound to dT_20_ and dT_20_ DNA templates. Both monomers and dimers bind dT_20_ (red) while only monomers bind dT_20_ (black). **(C)** Fluorescent anisotropy of a FAM labeled helicase template DNA (dT_20_ ssDNA tail with 18bp duplex) as a function of UvrD1 concentration. In the absence of DTT, both WT UvrD1 (red) and the 2B mutant (blue) displayed similar concentration dependencies of binding suggesting that they bind this template with similar affinity. **(D)** DNA unwinding traces as a function of UvrD1 concentration first treated with 1 mM DTT, followed by the addition of 2 mM H_2_O_2_. A control in the absence of oxidizing agent is shown for comparison (-H_2_O_2_).

The observation of DNA-bound monomers directly eliminates the possibility that monomers do not unwind because they do not bind DNA and suggests that some other property of the dimer is required for unwinding. This is consistent with other studies of UvrD-family enzymes as described in the Discussion.

At a constant redox potential established by the addition of 2 mM H_2_O_2_, the expected protein-concentration dependence of DNA unwinding is observed **(Fig. 5D)**. Here, the fraction of DNA unwound saturates at ~27% consistent with the fraction of dimers under these conditions **(Fig. 1D)** further suggesting that the monomers and dimers compete approximately equally for this DNA substrate under these conditions.

### Both UvrD1 monomers and dimers are single-stranded DNA translocases

As both monomers and dimers can bind to the DNA substrate **(Fig. 5)**, we considered our second hypothesis that postulated that monomers are unable to translocate along single-stranded DNA thus preventing helicase activity. To test this, we examined the translocation kinetics of UvrD1 on ssDNA. Since UvrD1 has a 3’ to 5’ DNA unwinding polarity, the kinetics of translocation were measured via stopped-flow assays (62, 63) by monitoring the arrival of UvrD1 at the 5’-end of a series of Cy3 5’-end-labeled oligodeoxythymidylate ssDNAs of different lengths *(L* = 20, 35, 45, 75, and 104 nucleotides) **(Fig. 6A)**. Arrival of a translocating protein at the 5’-end results in an enhanced Cy3 fluorescence and subsequent dissociation of UvrD1 leads to a return of the signal to baseline. A heparin concentration of 1 mg/ml was added to prevent rebinding of free UvrD1 to the ssDNA, ensuring single-round conditions **(Fig. S13A)**. In addition, negative controls in the absence of ATP showed no change in fluorescence **(Fig. S13A)** and experiments containing ATP and UvrD1 in the presence of 3’-labeled DNA showed only a decay in fluorescence consistent with translocation away from the dye **(Fig. S13B)**. In conditions favoring dimeric UvrD1, the presence of ATP resulted in the expected length-dependent peaks of fluorescence indicative of UvrD1 translocation in the 3’ to 5’ direction (62, 63) **(Fig. S14)**. However, monomeric UvrD1 generated either by the addition of DTT or the use of the 2B mutant also resulted in DNA-length-dependent changes in the fluorescence signal consistent with 3’ to 5’ ssDNA translocation **(Fig. 6B,C)**. Global analysis using an n-step sequential model **(Fig. S15)** (62, 64) produced better fits for the 2B mutant data possibly because the WT UvrD1 still contains trace amounts of dimer. (Methods and **Fig. S16**) However, fits of both data sets yielded consistent estimates of the macroscopic translocation rate *(mkt)* of 120 ± 5 nt/sec (WT+DTT) and 130 ± 10 nt/sec (2B-DTT) for monomeric UvrD1 which is very similar to the ssDNA translocation rate measured for *E. coli* UvrD monomers (62, 63). In addition, the estimated dissociation rate *(kd)* from the fitting analysis (4.1 ± 0.1 s^-1^) was on the same order of magnitude as that obtained by experimentally measures (8.1 ± 0.2 s^-1^) **(Fig. S16)**. Other fit parameters are listed in **Supplemental Table 4** (64).

**Figure 6:**
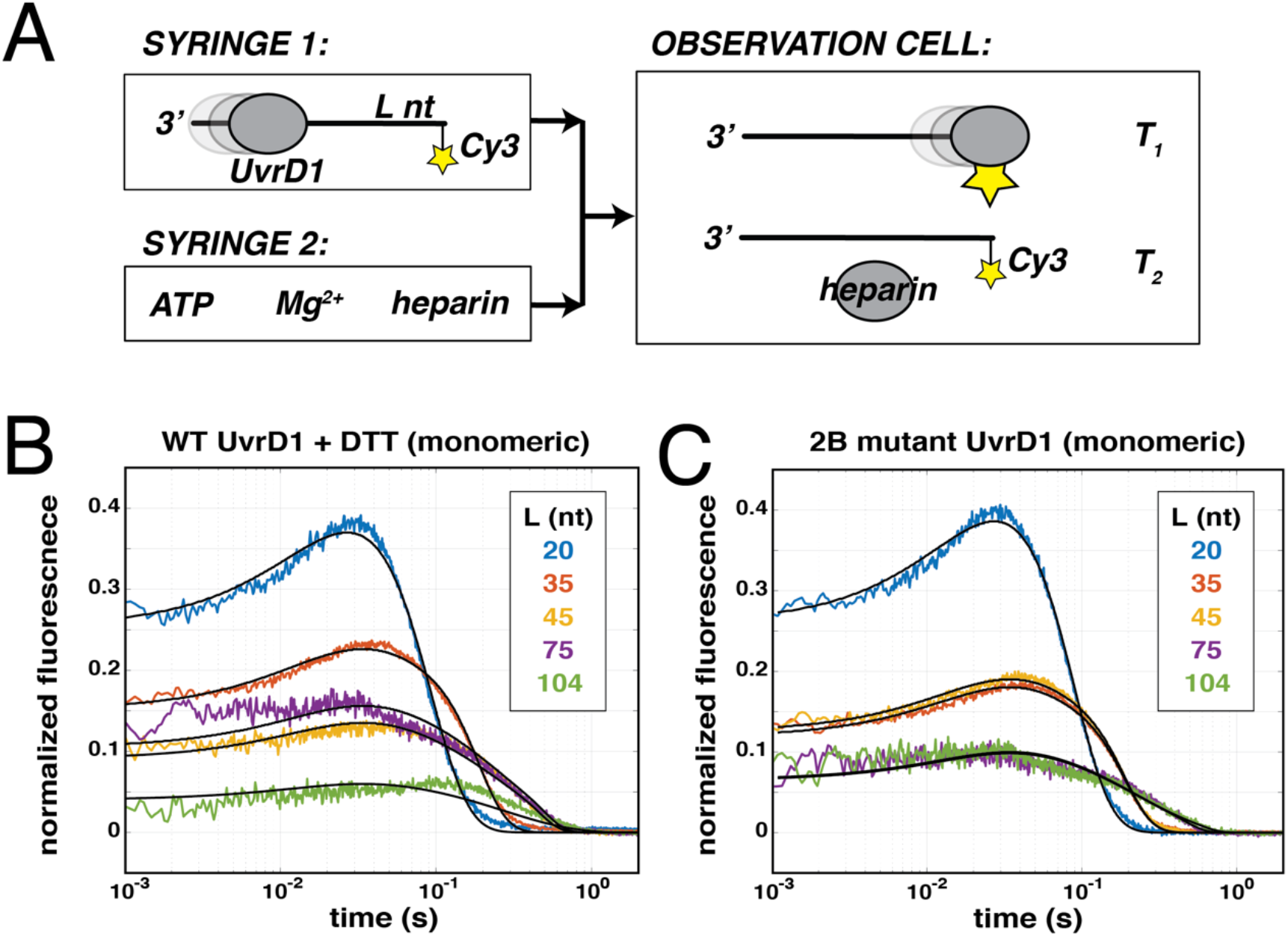
Monomeric UvrD1 translocates on single-stranded DNA. **(A)** The stopped-flow translocation assay. UvrD1 is allowed to equilibrate and bind to ssDNA templates of different lengths (L). These templates are labeled a the 5’ end with Cy3. The protein DNA solution is mixed with buffer containing ATP, Mg^2+^, and heparin to stimulate a single round of translocation. As the protein passes the label, the fluorescence increases (T1). As it dissociates, the fluorescence decreases, and free protein is bound by heparin (T2). **(B)** WT UvrD1 in the presence of 1 mM DTT produces traces consistent with translocation. The peak broadens and moves to longer times with increasing template lengths. **(C)** Monomeric UvrD1 generated by use of the 2B mutant, also shows clear peaks consistent with translocation. In both (B) and (C) experiments were conducted in Buffer A with 75 mM NaCl, where the traces were normalized to the average of 10 final plateau values observed in the raw data at each DNA length. Fits to an n-step sequential stepping model are shown in solid lines.

The time-courses exhibited under monomeric UvrD1 conditions were distinct from those collected under oxidative conditions suggesting that the monomer and dimer populations exhibit distinct translocation kinetics **(Fig. 6B,C, and Fig. S14)**. Fits using the percent fraction of monomeric and dimeric species and the translocation parameters obtained under monomeric conditions resulted in estimates for the ssDNA translocation properties of the UvrD1 dimer **(Supplemental Table 4)**. Although analysis of the mixed dimer/monomer population was challenging due to the presence of multiple species and kinetic phases, taken together thedata unequivocally show that both monomeric and dimeric UvrD1 translocate with 3’ to 5’ directionality along ssDNA.

### DNA unwinding activity is titrated by redox potential through dimer formation

We have shown that UvrD1 dimerization can be titrated via the addition of an oxidizing agent such as H_2_O_2_ **(Fig. 1D)**. Furthermore, these dimers are, in the absence of other activators, required for DNA unwinding **(Fig. 4)**. To determine the quantitative relationship between dimer fraction, DNA unwinding, and redox potential in millivolts (mV), we performed both sedimentation velocity and helicase assays using H_2_O_2_ to titrate redox potential. The 1A1B double mutant was used for these titrations to ensure that any effects stemmed directly from the cysteine in the 2B domain. We tested two different concentrations 1 μM and 2 μM 1A1B double mutant in the presence of 1 mM DTT which results in over 95% monomer **(Fig. 2D)** and shows no DNA unwinding **(Fig. 4C)**. We then titrated H_2_O_2_ from 0 – 5 mM which corresponds to redox potentials between −270 to 130 mV. As the redox potential became more positive (oxidizing), the fraction of DNA unwound increased **(Fig. 7, S17 purple)**. As in the case of the WT UvrD1 **(Fig. 1D)**, the fraction of UvrD1 1A1B double mutant dimer also increased with increasing H_2_O_2_ **(Fig. 7, orange)**. In fact, there is a quantitative correlation between the fraction of DNA unwound and the fraction of UvrD1 mutant dimer, consistent with the hypothesis that UvrD1 helicase activity is stimulated via increasing positive redox potentials found under oxidative conditions **(Fig. 7 and Fig. S17)**.

**Figure 7:**
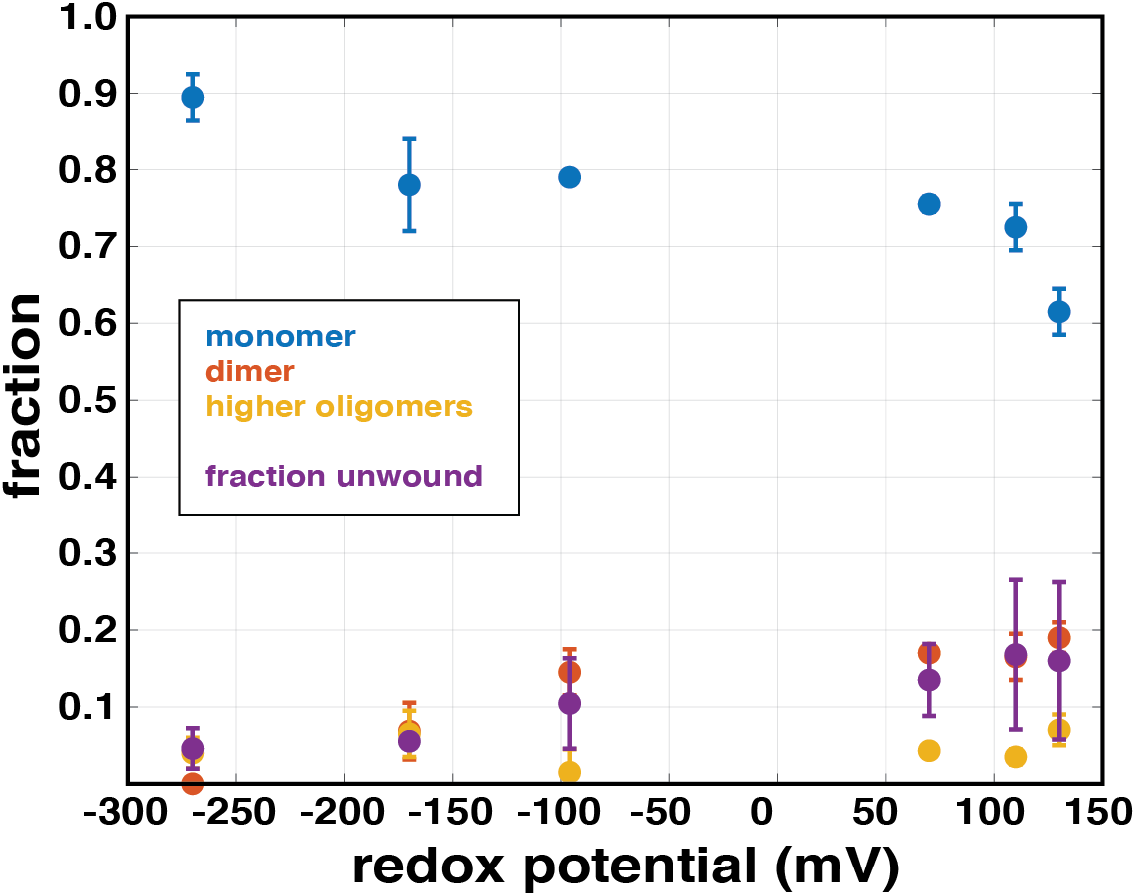
Unwinding activity correlates with dimer fraction and is titrated by redox potential. 1 μM 1A1B double mutant UvrD1 was dialyzed in Buffer A at 75 mM NaCl, treated with 1 mM DTT, and then incubated with varying concentrations of H_2_O_2_ (0-5 mM). Redox potential was measured and the samples were subject to AUC and used for DNA unwinding assays. The fraction of monomer present in different oligomeric states and fraction DNA unwound are plotted as a function of redox potential as follows: monomer (blue), dimer (orange), higher oligomers (yellow), and fraction DNA unwound (purple).

## Discussion

DNA helicases are enzymes that couple ATP binding and hydrolysis to translocation on ssDNA and unwinding of double-stranded DNA and are involved in critical pathways throughout nucleic acid metabolism. The SF1 superfamily of helicases is the largest group of known helicases and includes UvrD, Rep and PcrA (19, 20). These helicases consist of two RecA-like domains (1A and 2A) and two accessory sub-domains (1B and 2B) (19, 20). Studies of SF1 helicases have shown that in the absence of accessory factors or force exerted on the DNA, monomers possess ssDNA translocase activity, but not helicase activity; helicase activity requires at least a dimeric form of the enzyme. (21,22, 27, 28, 30, 32, 33, 62-64)Yet, the dimerization interface is not known as crystal structures of UvrD and PcrA only reveal a monomer bound to DNA (65–68).

Previous studies of *Mtb* UvrD1 concluded that it is monomeric as shown by size exclusion chromatography (SEC) and equilibrium sedimentation and is capable of unwinding dsDNA in this form (45, 48). However, in one case, SEC and equilibrium sedimentation were performed in the presence of the reducing agent DTT, whereas DNA unwinding assays were performed in its absence (48). In the other case, helicase activity of UvrD1 was attributed to monomers and was dependent on its binding partner, Ku (45). When we purified *Mtb* UvrD1 without reducing agent it eluted as two peaks on SEC corresponding to the expected molecular weights of both monomer and dimer **(Fig. 1A)**. AUC experiments show that, while salt concentration had almost no effect on the amount of the dimeric form, the addition of DTT results in a dramatic shift to a single monomeric peak suggesting that *Mtb* UvrD1 undergoes a cysteine dependent dimerization **(Fig. 1)**. We also observe a higher oligomeric species increasing in a concentration dependent manner in sedimentation velocity experiments performed under oxidizing conditions. We do not observe *Mtb* UvrD1 tetramer binding to the tailed construct used in our unwinding assays **(Fig. 5A)**, but we cannot eliminate the possibility that the tetrameric form could bind to DNA possessing longer singlestranded tails.

We identified a critical cysteine residue in the 2B domain of UvrD1 (C451) that is required for UvrD1 dimerization **(Fig. 2)**. More specifically, the C451A mutation abrogated both dimerization and helicase activity pointing to the existence of a 2B-2B disulfide bonded dimer. The 2B domain of UvrD-like helicases has been described as an auto-inhibitory domain (28, 32) that can adopt a range of rotational conformational states relative to the rest of the protein (28, 32, 66). The different 2B domain conformational states of the monomer are influenced by salt concentration, DNA binding, enzyme dimerization and the binding of accessory protein factors (26, 34, 69, 70). For example, crystal structures of Rep bound to ssDNA showed Rep monomer bound to DNA in two different conformations (66). In one of the conformations the 2B domain is in an “open” conformation, while in the other, the 2B domain is reoriented by a 130° swivel motion around a hinge region to contact the 1B domain. This swiveling motion closes the binding groove located between 1A, 1B, and 2A domains around the DNA template. Consistently, single molecule and ensemble FRET studies have also shown that the 2B domain of a monomer can be in closed or open conformations (31, 34, 69–71). Furthermore, removing the 2B domain from Rep causes the Rep monomer to gain helicase activity (32). These observations and others have led to the hypothesis that the 2B domain serves as the interface between subunits within the functional Rep and UvrD dimers (19, 28, 30, 72).

The formation of a 2B-2B disulfide bond in the case of *Mtb* UvrD1 fits well in a model where 2B-2B driven dimerization results in an active helicase conformation. Further support for this idea can be found in the specific location of C451. Mutations to 2B domain threonine residues such as T426 of *B. stearothermophilus* PcrA and T422 of *E. coli* UvrD disrupt helicase activity (73). This threonine residue is conserved in non-Actinobacteria UvrDs, is absent from UvrD-like proteins containing the cysteine residue described here and is located 2 amino acids upstream to the C451 of *Mtb* UvrD1. The similar location of these residues suggests that UvrD-like helicases share a common 2B-2B dimerization interface and that mutating this threonine destabilizes the dimeric form of the enzyme. Interestingly, the 2B domains of the SF1 family members RecB and RecC interact in a RecBCD helicase complex (74). We aligned the UvrD1 2B domain sequence with the 2B domain sequences of *E. coli* RecB and RecC which lack a 2B cysteine equivalent to C451. When we mapped residues that aligned close to the 2B cysteine onto the structure of RecBCD (PDB:5LD2) (74, 75), we observed that these regions in RecB and RecC are only 10-15Å apart. Again, this is consistent with our identification of the region surrounding the 2B cysteine of UvrD1 as an interface for 2B-2B domain-based activation of UvrD1-like helicases.

As noted, the 2B cysteine we have identified in *Mtb* UvrD1 appears unique to certain classes of Actinobacteria **(Fig. 3)**. The 2B domain of *E. coli* UvrD does contain a cysteine residue at a different location (C441). However, experiments with *E. coli* UvrD showed no effect on oligomeric state with changes in redox potential **(Fig. S7)**.

Another pathway for stimulation of unwinding by UvrD-family enzymes requires the binding of activating partners. The mismatch-repair protein MutL can activate the monomer helicase activity of *E. coli* UvrD as well as stimulate the activity of UvrD dimers and activation is accompanied by a change in the rotational state of the 2B domain (33). This interaction also leads to enhanced processivity which enables UvrD to unwind longer stretches of DNA when functioning with MutL (32). Similarly, the accessory factor PriC can activate the Rep monomer helicase and stimulate the Rep dimer helicase (76). By analogy, the NHEJ factor, *Mtb* Ku, has been reported to bind to the C-terminal region of UvrD1 (45) suggesting a role for UvrD1 in double-strand-break repair (DSB). As the results from our study show that UvrD1 can exist as a monomer or dimer depending on the redox potential, we are currently investigating whether Ku can activate the UvrD1 monomer helicase and/or stimulate dimer activity and whether it does this via modulation of the 2B domain conformation.

*Mtb* actually has two UvrD-family members in its genome, UvrD1 and UvrD2, with UvrD1 being the homologue to *E. coli* UvrD (45–47, 77) and UvrD2 consisting of an N-terminal SF1 helicase motor linked to an HRDC (helicase and Rnase D C-terminal) domain and tetracysteine motif domains (44, 78). The cysteines within the tetracysteine motif bind zinc and the domain is required for helicase activity of UvrD2 *in vitro* (44). However, in this case the activity appears to be dependent on the presence of the domain and not the presence of the individual cysteines themselves. Interestingly, while WT UvrD2 does not show a dependence on Ku, a truncated construct lacking the tetracysteine domain can be activated by Ku (44). Therefore, while both UvrD1 and UvrD2 utilize cysteine residues for helicase activity, they do so via distinct biochemical mechanisms.

During infection, *Mtb* resides within alveolar macrophages and neutrophils where it is exposed to reactive oxygen species (ROI) and reactive nitrogen intermediates (RNI) that cause DNA damage (49, 79, 80). *Mtb* lacks mismatch repair (81) and the response of *Mtb* to these insults likely involves NER pathways (79). During NER, UvrD1 unwinds the damaged DNA strand to remove bulky lesions that have been recognized by UvrA/UvrB and excised by UvrC (2). Analysis of gene expression data have shown UvrA to be highly expressed during oxidative stress (82). In addition, both *uvrD1/uvrB* and *uvrD1/uvrA* double mutants in *M. smegmatis* have been shown to be more sensitive to tertiary butyl hydroperoxide and acidified nitrite than wild-type strains (46, 83). All of these observations suggest a role of NER enzymes, including UvrD1, during oxidative stress in *Mtb*. This is distinct from *E. coli,* in which repair of RNI and ROI-induced DNA damage is accomplished by base-excision repair and homologous recombination (84). Thus, the redox dependent dimerization of UvrD1 we report here may represent an important mechanism in *Mtb* underlying the repair of oxidative-dependent DNA damage during infection.

## Materials and Methods

### Cloning, overexpression, and purification of *Mycobacterium tuberculosis* UvrD1

*Mtb* UvrD1 (Rv0949) from H37Rv was cloned in the expression vector pET with SUMO-His tag at the N- terminus and Kannamycin resistance. It was PCR amplified with BamH1 at the 5’ and HindIII at the 3’end. UvrD1 2B domain cysteine to alanine substitution and 1A1B double cysteine to threonine substitution mutations were introduced into the *Mtb* UvrD1 plasmid by PCR amplification using primers and site directed mutagenesis kit Agilent product 200521. Sequences for all primers and a list of plasmids can be found in **Supplemental Table 5**. The inserts of all UvrD1 plasmids were sequenced to exclude the acquisition of unwanted coding changes during amplification or cloning. The pET-*Mtb* UvrD1 plasmids were transformed into *Escherichia coli* BL21(DE3). Cultures (3 L) were grown at 37 °C in a Luria-Bertani medium containing 50 mg/ml kanamycin until the *A*_600_ reached ~0.5. The cultures were chilled on ice for about 1 hour, and the expression of recombinant protein was induced around 0.55 OD with 0.25 mM isopropyl-β-D-thiogalactopyranoside (IPTG), followed by incubation at 16 °C for 16 h with constant shaking. The cells were harvested by centrifugation, and the pellets were either stored at −80 °C or used for subsequent procedures that were performed at 4 °C. The bacterial cells from the 3-liter culture were resuspended in 50-75 ml of lysis buffer (50 mM Tris-HCl, pH 7.5, 0.25 M NaCl, 10% sucrose). Lysozyme and Triton X-100 were added to final concentrations of 1 mg/ml and 0.1%, respectively. At the time of lysis, a complete EDTA-free protease inhibitor cocktail (Sigma: product 118735800) was added to the lysate. Next the lysates were sonicated and insoluble material was removed by centrifugation at 14K for 45 minutes. The soluble extracts were applied to 2-ml columns of nickel-nitrilotriacetic acid-agarose (Ni-NTA) (QIAGEN catalog no. 30210) that had been equilibrated with lysis buffer without protease inhibitors. The columns were washed with 10X column volume of wash buffer (50 mM Tris-HCl, pH 8.0, 0.25 M NaCl, 0.05% Triton X-100, 10% glycerol) and then eluted stepwise with wash buffer containing 50, 100, 200, 500, and 1000 mM imidazole. The polypeptide compositions of the column fractions were monitored by SDS-PAGE. The his-SUMO-tagged UvrD1 polypeptides were recovered predominantly in the 100- and 200-mM imidazole eluates. Fractions containing the UvrD1 protein were pooled and His-tagged Ulp1 protease was added (at a ratio of 1:500 wt/wt protease per protein) and dialyzed against dialysis buffer (50 mM Tris-HCl, pH 8.0, 1 mM EDTA, 0.1% Triton X-100, 10% glycerol) containing 150 mM NaCl overnight. The SUMO-cut UvrD1 was then incubated with Ni-NTA agarose for about 3 hours and the untagged UvrD1 (cleaved) was recovered in the flow-through and wash fractions. After pooling and concentrating the fractions by VIVASPIN centrifugal filters (30 kDa cutoff: product Sartorius VS2021), the protein was loaded on heparin HiTRAP column (Cytiva: product 17040701) (5 ml x 2) pre-equilibrated with dialysis buffer containing 150mM NaCl. Upon running a linear gradient from 200 to 800 mM NaCl, the protein eluted at approximately 400 mM NaCl. The fractions with a single band on a reducing SDS-PAGE corresponding to the molecular weight of a monomer were pooled together and concentrated to load on the S300 sizing column (HiPrep 16/60 Sephacryl S-300 HR column,Cytvia, product 17116701) in buffer 150mM NaCl, 10% glycerol and TRIS pH 8.0 at 25 °C without DTT. The peak fractions were pooled and stored in −80 °C at concentrations about 15-20 μM. Both the 2B mutant and 1A1B double mutant were overexpressed and purified in a manner similar to the wild-type UvrD1 protein.

### Homology Modeling of *Mtb* UvrD1

The predicted structure of *Mtb* UvrD1 was obtained using PHYRE2 (85) by submitting the amino acid sequence of *Mtb* UvrD1. The PDB reference used for modeling open structure of *Mtb* UvrD1 was *E. coli* UvrD was 3LFU and for modeling closed structure is *B. stearothermophilus* PcrA helicase complex 3PJR and methods for SASA calculations done in Chimera (53).

### Phylogeny analysis of *Mtb* UvrD1

The protein sequence of *Mtb* UvrD1 obtained from uniprot (https://www.uniprot.org/uniprot/P9WMQ1.fasta) was used to Blast against various genus and species of classes of Phylum Actinobacteria (https://blast.ncbi.nlm.nih.gov/Blast.cgi). Two-three genus of orders in which the cysteine in 2B domain is conserved were chosen and aligned in Clustal omega (86) and the phylogenetic tree was plotted using NJ plot (87).

### Analytical Size Exclusion Chromatography

1 mL of *Mtb* UvrD1 (30 μM) was injected in a S300 gel filtration column at a flow rate of 0.25 mL/min, monitoring absorbance at 280 nm for a measure of the elution volume *(V)* in buffer 150mM NaCl, 10% glycerol and TRIS pH 8.0. The value of *V_e_/V_o_* was interpolated using the generated standard curve (Bio-Rad gel filtration standard; product #1511901) to yield the estimated molecular weight of *Mtb* UvrD1 monomer 90 kDa and dimer 170 kDa fractions.

### DNA Substrates

Single stranded DNA which are either labeled with Cy5, Cy3, FAM or BHQ2 were ordered from IDT. For annealing of the oligos the Cy5 labeled at 5’end of the single stranded DNA was mixed with an equimolar concentration of unlabeled complementary strand or complementary strand labeled with BHQ2 in 10 mM Tris pH 8.0, 50 mM NaCl, followed by heating to 95 °C for five minutes and slow cooling to room temperature.

### Synthesis of poly-dT

The homodeoxypolynucleotide, poly-dT substrate was used to measure dissociation rate from internal sites of ssDNA. Since the poly-dT from commercial sources is polydisperse, we prepared samples using enzyme terminal deoxynucleotidyl transferase (TdTase) from calf thymus gland to catalyze polymerization of deoxynucleotide triphosphate into poly-dT of more well-defined lengths (88). The protocol includes mixing dT(100) with potassium cacodylate buffer, potassium chloride, Cobalt chloride, Inorganic pyrophosphatase (ThermoFisher Scientific #EF0221), deoxythymidine-5’-triphosphate dTTP (ThermoFisher Scientific #R0171) and terminal deoxynucleotidyl transferase (TdTase ThermoFisher Scientific #10533065). The reaction was kept at room temperature for 3-4 days and poly-dT was purified using phenol chloroform extraction and suspending the air-dried pellet in water. The weight average length of the poly-dT was determined by measuring the weight average sedimentation coefficient by boundary sedimentation velocity experiments using AUC. The weight average length determined from this method is 964 nucleotides and was determined using method from the measured weight average sedimentation coefficient on poly-U (89).

### Analytical ultracentrifugation

The analytical ultracentrifugation sedimentation velocity experiments were performed using a Proteome Lab XL-A analytical ultracentrifuge equipped with an An50Ti rotor (Beckman Coulter, Fullerton, CA). The sample (380 μl) and buffer (410 μl) were loaded into each sector of an Epon charcoal-filled two-sector centerpiece. All experiments were performed at 25 °C and 42,000 rpm. Absorbance data were collected by scanning the sample cells at intervals of 0.003 cm, monitoring either at 230 nm or 280 nm depending on protein concentration to maintain an absorbance signal between 0.1 and 1. Both the DNA and protein samples were dialyzed in buffer 75 mM NaCl, 20% glycerol and 10 mM TRIS pH 8.0 except the salt titration where the protein was dialyzed in different salts at 20% glycerol at 10mM TRIS pH 8.0.

Continuous sedimentation coefficient distributions, *c*(*s*), were calculated using SEDFIT(6), truncating the fit at 7.0 radial position to avoid contributions of glycerol buildup (90). This analysis yielded individual sedimentation coefficients for each monomer, dimer, and tetramer species as well as a weighted average frictional coefficient *(f/f_o_)* for the entire distribution (Table S1). Calculated sedimentation coefficients were converted to 20°C water conditions (*s_20,w_*) according to:

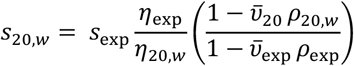

where *ρ_20w_* and *η_20w_* are density and viscosity of water at 20 °C, *ρ*_exp_ and *η*_exp_ are density and viscosity of the buffer at the experimental temperature of 25 °C, and 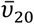 and 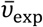 are partial specific volumes of the protein at 20 °C and at 25 °C. Buffer densities, (*ρ*_exp_) and viscosities (*η*_exp_) were calculated from buffer composition using SEDNTERP (91). Partial specific volumes 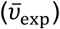 for *Mtb* UvrD1 and point mutations were calculated in SEDNTERP using the amino acid composition. Integration of the entire c(s) distribution vs. the integration of an individual sedimentation species was performed and used to calculate the population fraction (92).

For AUC experiments done in the presence of Cy5 labeled DNA, the absorbance signal was collected by scanning the sample cells at 650 nm. Partial specific volumes 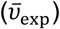 for labeled DNA and the UvrD1-DNA complex were calculated according to:

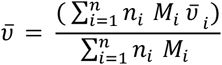

For AUC experiments conducted at different redox potentials, the protein was first dialyzed in Buffer A with 75 mM NaCl and then 1 mM DTT was added at a respective concentration and then titrated with a range of H_2_O_2_ from 0.5 to 2 mM. Redox potential was measured via Metler Toledo Redox micro electrode product #UX-35902-33. After H_2_O_2_ treatment protein was incubated at room temperature for two hours before performing AUC.

### Stopped-flow double-stranded (ds) DNA unwinding assay

All stopped-flow experiments were carried out at 25°C using an Applied Photo physics instrument SX-20, total shot volume 100 μl, dead time 1 ms. All experiments were carried out in Buffer with TRIS pH 8.0, 75 mM NaCl, and 20% glycerol in the absence or presence of 1 mM DTT. Cy5 fluorophore was excited using 625 nm LED (Applied Photo physics Ltd., Leatherhead, UK) and its fluorescence emission was monitored at wavelengths >665 nm using a long-pass filter (Newport Optics). The traces represent the average of 5 independent shots and at least two different protein purifications. In this assay, a double-stranded 18-basepair DNA with a T20/T10 tail with Cy5 fluorophore is attached to the long strand and the black hole quencher (BHQ_2) attached to the short strand (**Supplemental Table 6**). The concentrations mentioned are the final after mixing the contents of both syringes. UvrD1 (200 nM) is incubated with 2 nM DNA in one syringe which is then rapidly mixed with the contents in another syringe that consists of (TRAP a complementary DNA strand in excess protein (25X, 5 μM), Mg^+2^ (5 mM), and ATP (1 mM). The concentrations of ATP and Mg^+2^ used were determined to be optimal for *Mtb* UvrD1 unwinding assays (48). DNA strand separation is accompanied by an increase in the fluorescence signal. The unwinding signal is normalized to the signal from positive and negative control to get fraction of DNA unwound. The positive control is 2 nM double stranded DNA with BHQ2 on the short and Cy5 on the long strand that is denatured in the presence of TRAP and 200 nM UvrD1 is added at room temperature to get maximum fluorescence signal. The negative control is the average fluorescence value recorded for the fully annealed DNA (2 nM) with UvrD1 (200 nM) shot against buffer alone. To get DNA unwinding with a change in redox potential the protein was dialyzed treated with DTT and titrated with H_2_O_2_ in similar way as done for AUC experiments and incubated with DNA. This protein DNA mix was then used for unwinding experiments.

### Stopped flow single stranded (ss) DNA translocation assays

The kinetics of UvrD1 monomer and dimer translocation was examined in a stopped-flow experiment by monitoring the arrival of UvrD1 of a series of oligodeoxythymidylates length (L= 20, 35, 45, 64, 75, 94, and 104) nucleotides labeled at the 5’-end with Cy3 **(Supplemental Table 6)** (62). Cy3 fluorescence was excited using a 535 nm LED with a 550 nm short-pass cut-off filter and emission was monitored at >570 nm using a long-pass filter (Newport Optics). UvrD1 was pre-incubated with ssDNA in one syringe and reactions were initiated by 1:1 mixing with 1 mM ATP, 5 mM MgCl_2_ and heparin at a concentration of 1 mg/ml to prevent rebinding of UvrD1 (50 nM) to DNA (100 nM). Excess DNA to UvrD1 ratio was used to prevent binding of more than one UvrD1 monomer on DNA (62). Global analysis of time courses using the n-step sequential model was done to calculate various translocation parameters. For making heparin solution to be used as a TRAP for protein ensuring single round conditions, heparin sodium salt (porcine intestinal mucosa, Millipore Sigma #H3393) was dialyzed into Buffer A plus 75 mM NaCl and concentrations were determined by an Azure A standard curve (93).

### Tryptophan fluorescence-based dissociation kinetics

Dissociation kinetics of UvrD1 were monitored by the increase in UvrD1 tryptophan fluorescence, excited using a 290 nm LED (Applied Photophysics Ltd., Leatherhead, UK) and monitoring emission at >305 nm using a long-pass filter (check if this is from Newport or AP). The observed dissociation rate from internal ssDNA sites for UvrD1 monomer (with 1 mM DTT) and dimer (no DTT) was measured using (dT)100 and poly(dT) oligos (average length of 964 nucleotides; see synthesis of poly-dT section), respectively. In one syringe UvrD1 (100 nM) was added with DNA (50 nM) (concentrations listed are after equal volume mixing). In another syringe ATP, MgCl_2_ and heparin were added at the same concentrations as translocation assays and the observed dissociation kinetic traces were best fit to a single exponential using ProData Viewer (Applied Photophysics). All experiments were performed at 25°C in Buffer A with 75 mM NaCl and represent the average of 5 independent shots (88).

### ssDNA translocation and unwinding fitting analysis

The translocation and the unwinding data was fit using the code from https://github.com/ordabayev/global-fit and unwinding and translocation rates are calculated. Globalfit is a wrapper around lmfit https://lmfit.github.io/lmfit-py/ providing an interface for multiple curves fitting with global parameters. Python 3 is installed via Anaconda along with modules like numpy, scipy, matpotlib, lmfit, emcee, corner, os and pandas and then globalfit model is used to fit the data for unwinding using n-step unwinding model and translocation using a two-step dissociation model (64).

### Steady-State Anisotropy Measurements

All fluorescence titrations were performed using a spectrofluorometer (ISS, Champaign, IL) equipped with Glan-Thompson polarizers. Measurements of the anisotropy and total fluorescence intensity of FAM-labeled double stranded DNA 18bp with T20 single stranded tail were recorded using excitation and emission wavelengths of 490 and 522 nm, respectively, using

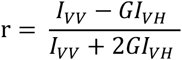

where *I_TOT_* = *I_VV_* + *2GI_VH_* and *G* is the *G* factor (94). The recorded value of G factor remained between 0.85 to .9 throughout the titrations. Titrations were performed using a Starna cells cuvette catalog number 16.100F-Q-10/Z15 with dimensions 12.5 x 12.5 x 45 mm and pathlength 1cm. The protein was mixed 3-4 times with DNA during titrations and let it sit for 5 minutes before recording anisotropy values. The total volume of added protein was 30% of the initial volume and as a control the dilution with buffer of up-to 30% for DNA alone sample did not change the anisotropy value of the DNA. All titrations were conducted in Buffer A at 25 °C in 75 mM NaCl. The data represents average from three independent experiments for WT-DTT and 2 independent experiments for the 2B mutant.

## Acknowledgments

This work was supported by NIH R01 GM134362 to EAG, NIH R35 GM136632 to TML, and by NIH T32 AI007172 to AC. The content is solely the responsibility of the authors and does not necessarily represent the official views of the National Institutes of Health.

