## Supplemental Information for "The *Mycobacterium tuberculosis* DNA-repair helicase UvrD1 is activated by redox-dependent dimerization via a 2B domain cysteine conserved in other Actinobacteria"

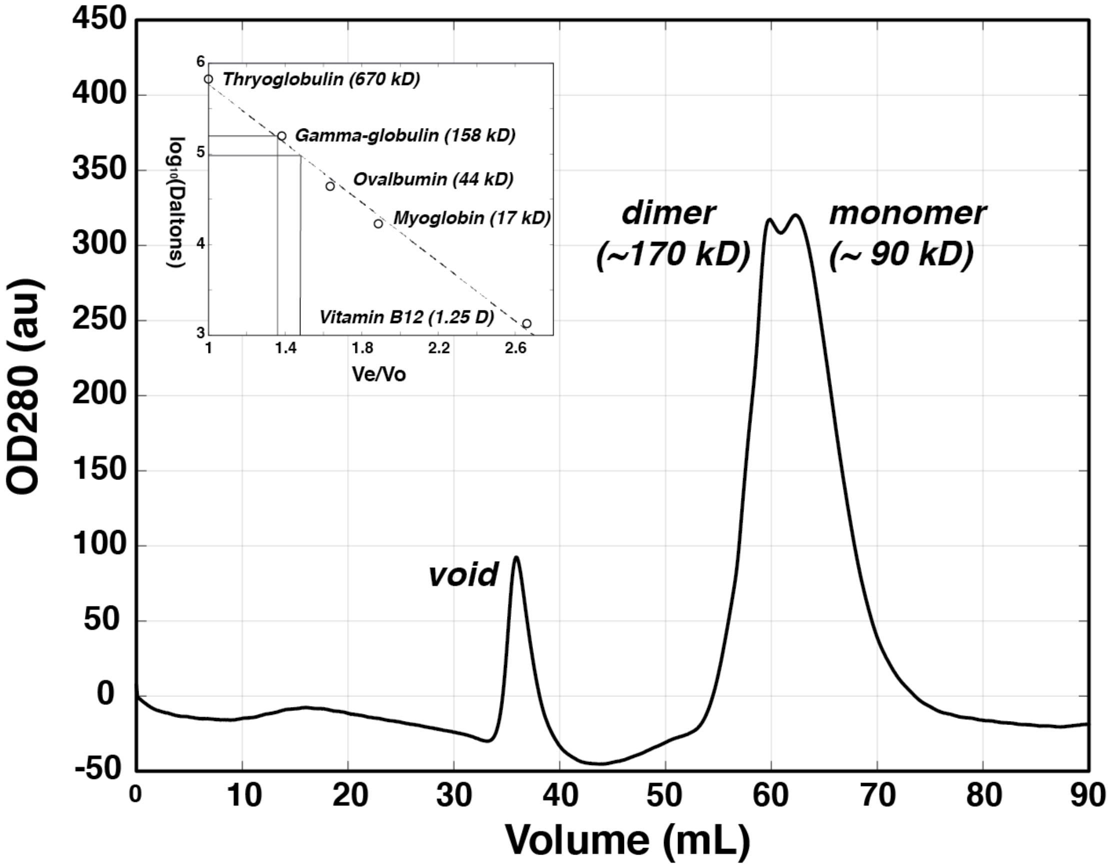


**Supplemental Figure 1: Size exclusion chromatogram indicating the presence of multiple oligomeric forms of UvrD1.** A S300 size exclusion column was run in 150 mM NaCl, 10% (v/v) glycerol, 10mM TRIS pH 8.0 at 25 $^{o}$C in the absence of DTT at 4 $^{o}$C. Standards run on the same column were used to estimate the molecular weight of the two peaks (see Methods) (insert). The first peak is consistent with the presence of UvrD1 dimers (170,000 Da measured compared to 170,000 Da theoretical) while the second is consistent with the presence of UvrD1 monomers (90,000 Da compared to 85,000Da theoretical). We note here that depending on the resolution of individual sizing runs, this doublet peak is not observed upon purification; instead, a long tail at larger molecular weights is observed.

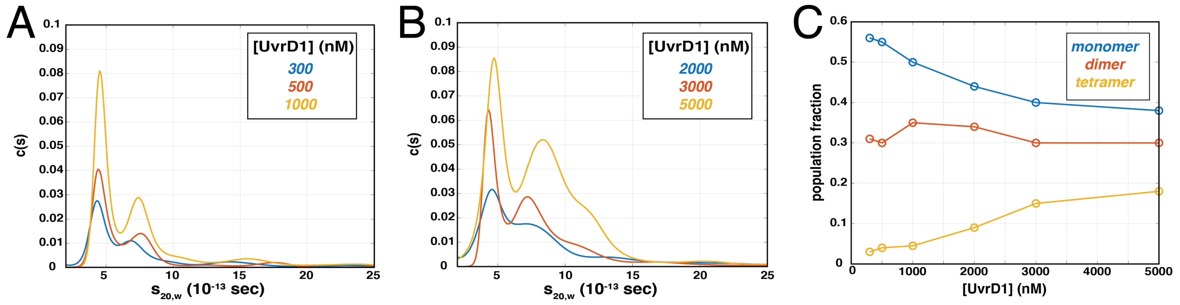


**Supplemental Figure 2: AUC sedimentation velocity c(s) distributions of UvrD1 as a function of protein concentration.** All experiments were performed in Buffer A with 75 mM NaCl and 1mM DTT at respective concentrations and treated with 2mM $H_2O_2$. **(A)** 300 nM (blue), 500 nM (red), and 1000 nM (yellow) runs were monitored at 230 nm. **(B)** 2000 nM (blue), 3000 nM (red), and 5000 nM (yellow) runs were monitored at 280 nm. **(C)** The population fraction of monomer (blue), dimer (red), and tetramer (yellow) were calculated by integrating the data from A and B and are plotted as a function of protein concentration.

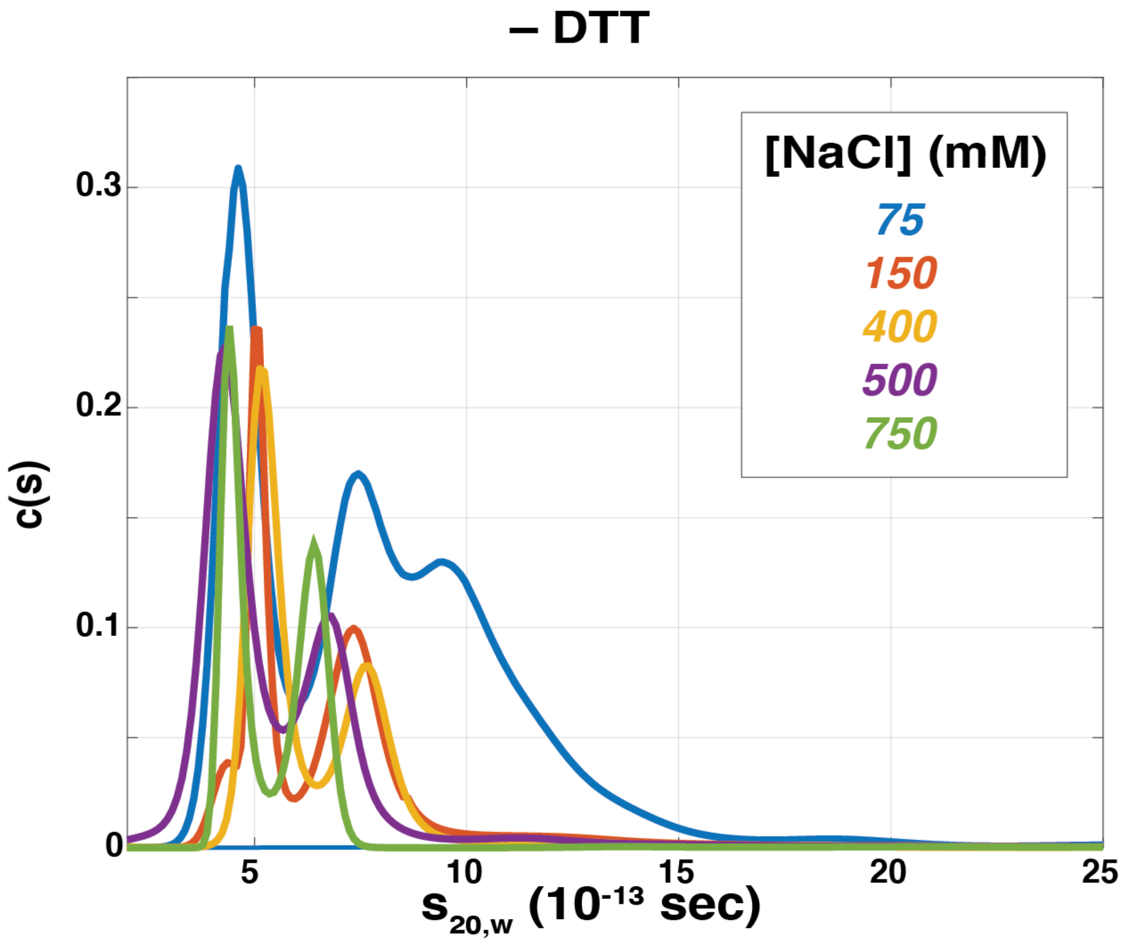


**Supplemental Figure 3: AUC sedimentation velocity c(s) distributions at different NaCl concentrations.** Sedimentation velocity traces used to measure fractions of monomer, dimer, and higher order oligomers presented in the main text (Fig. 1B). Experiments were performed at a constant 2.5 µM protein concentration in Buffer A (TRIS pH 8.0 at 25 C, 20% glycerol) without DTT at the NaCl concentration indicated in the legend. All traces are averages of two separate experiments and were monitored at 280 nm except for 75 mM NaCl which was monitored at 230 nm. Therefore, the amplitudes of 75 mM condition should not be compared to the other traces.

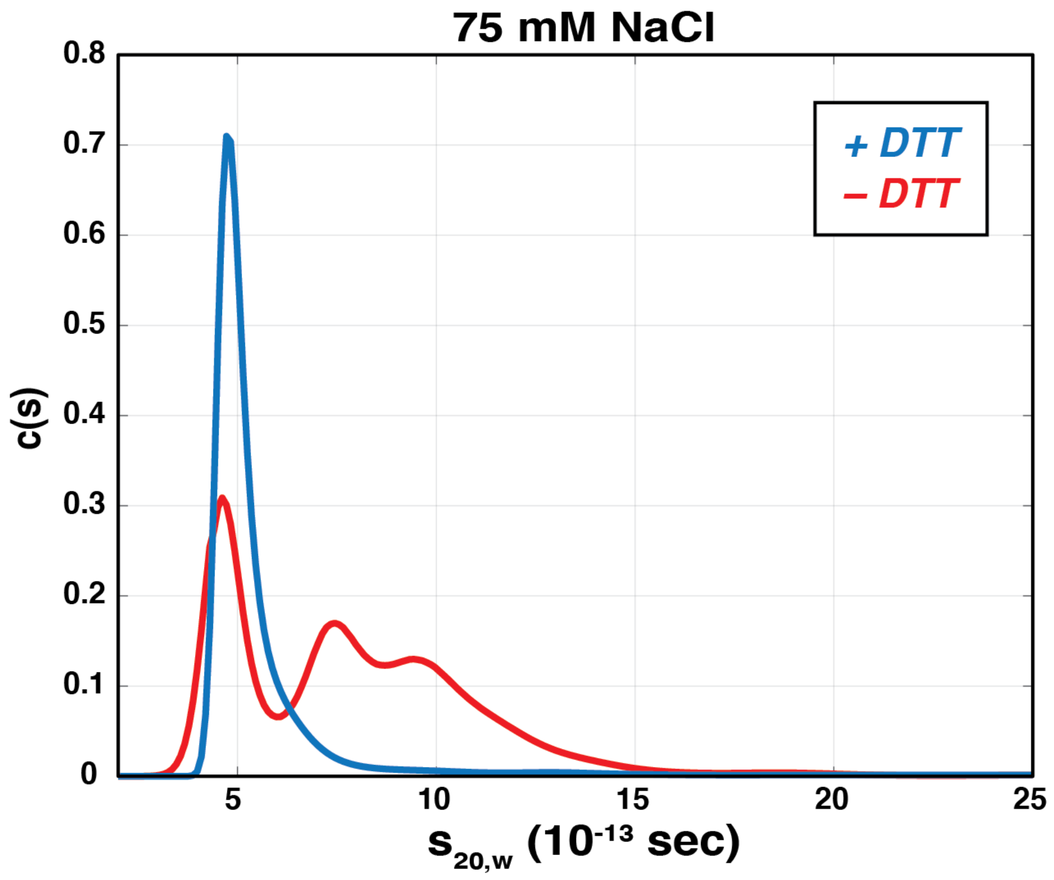


**Supplemental Figure 4: DTT dependence of dimerization at 75 mM NaCl. AUC sedimentation velocity c(s) distributions** display a similar effect of DTT in Buffer A with 75 mM NaCl compared to the results in the main text at 400 mM NaCl (Fig. 1C). Each trace is an average of two separate experiments monitored at 230 nm at 2.5 µM protein concentration.

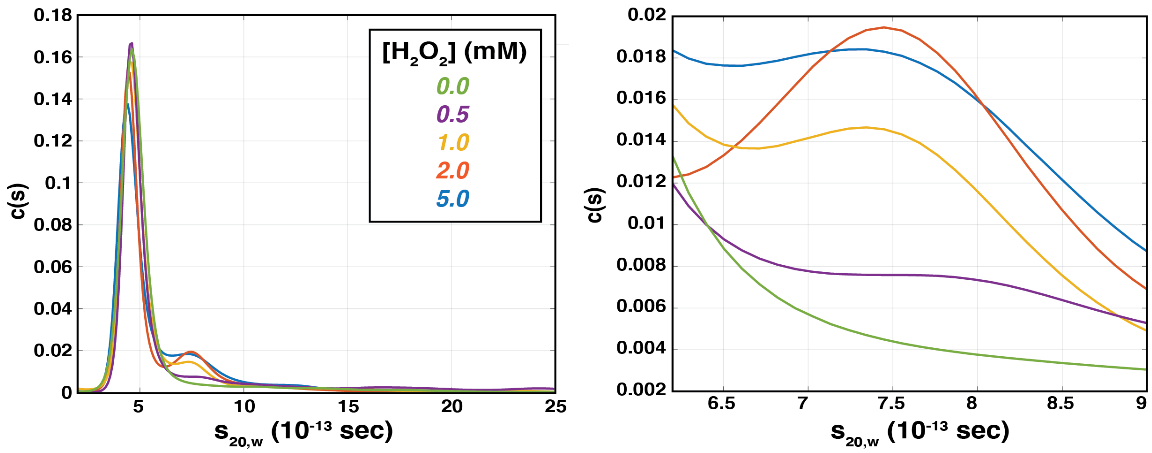


**Supplemental Figure 5: AUC sedimentation velocity c(s) distributions at different hydrogen peroxide concentrations.** 2.5 µM UvrD1 was dialyzed in Buffer A with 75 mM NaCl and 1 mM DTT and then subjected to $H_2O_2$ addition (see Methods). The dimer fraction was found to increase as a function of $H_2O_2$ concentration (quantified in Fig. 1D). (Right) A zoom into the region representative of the dimer is shown for clarity. Each trace is an average of two separate experiments, monitored at 280 nm.

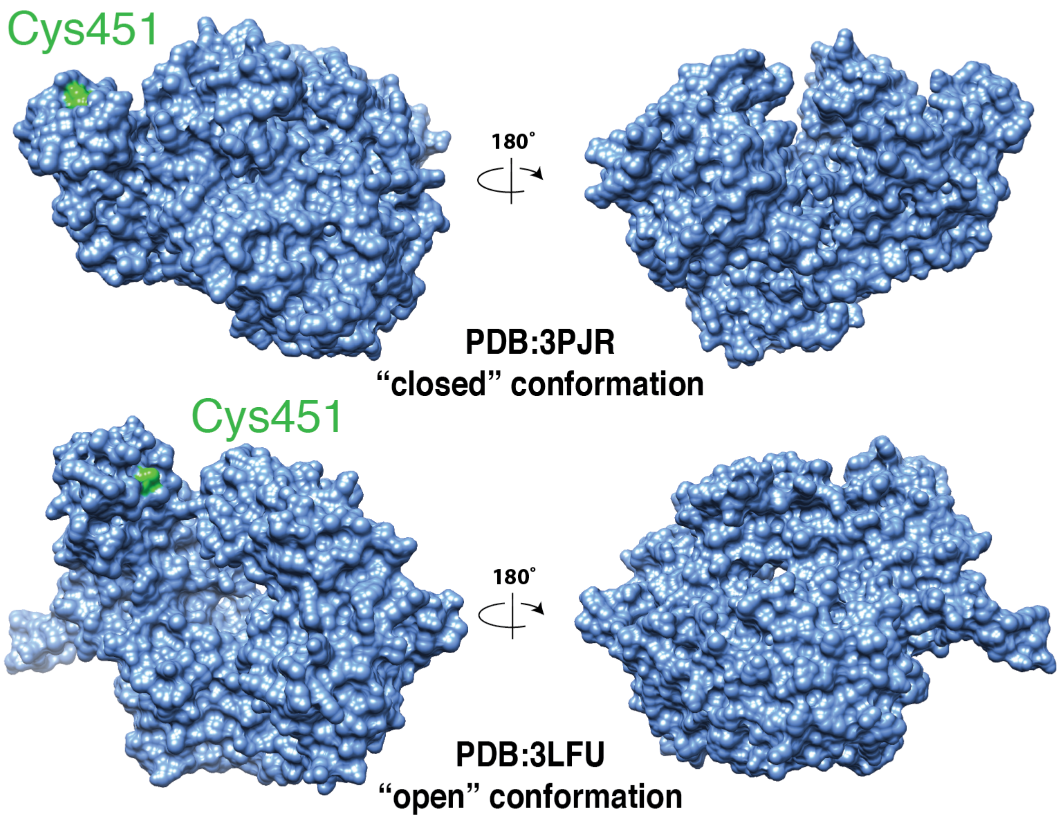


**Supplemental Figure 6: (A)** A surface view of the threaded *Mtb* UvrD1 structure based on the *G. stearothermophilus* UvrD “closed” structure (PDB:3PJR). **(B)** A surface view of the threaded *Mtb* UvrD1 structure based on the *E. coli* “open” UvrD structure (PDB:3LFU). In both cases, of the three total cysteines in the *Mtb* UvrD1 sequence, only C451 in the 2B domain is surfaced exposed (green).

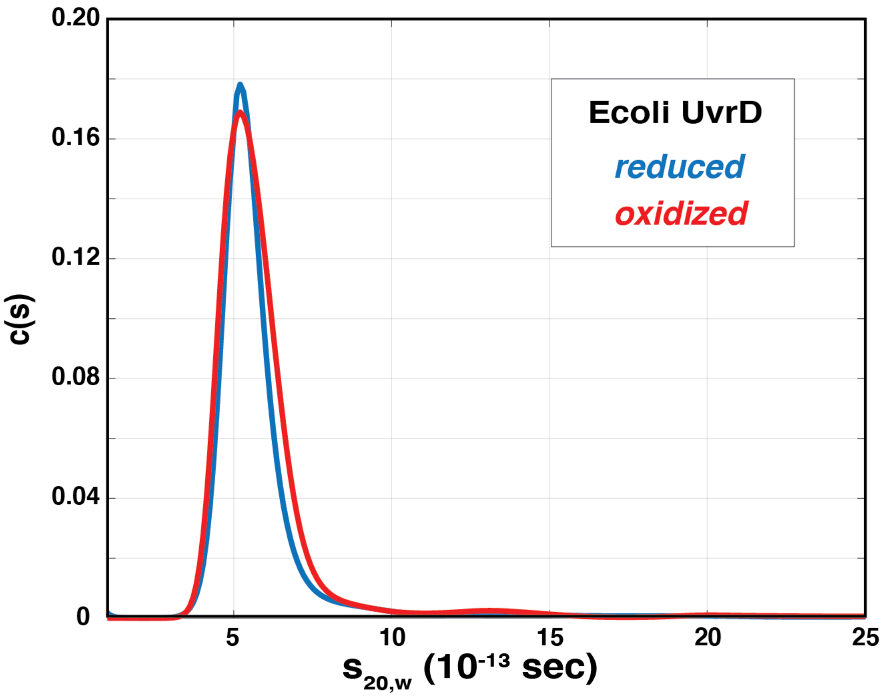


**Supplemental Figure 7: *E. coli* UvrD does not dimerize in a redox-dependent manner.** Sedimentation velocity was run with *E. coli* UvrD in both reducing and oxidizing conditions. Briefly, 4 µM UvrD stored in 200 mM NaCl, 50% (v/v) glycerol,10 mM TRIS pH 8.3 25 $^{o}$C and 25 mM beta-mercaptoethanol was first dialyzed in Buffer A with 75 mM NaCl and 2 µM protein was incubated with either 1 mM DTT (blue) or 1 mM DTT with 2 mM $H_2O_2$ (red). The resulting c(s) distributions indicated a single species observed in both redox conditions, suggesting that *E. coli* UvrD does not dimerize in response to oxidative conditions.

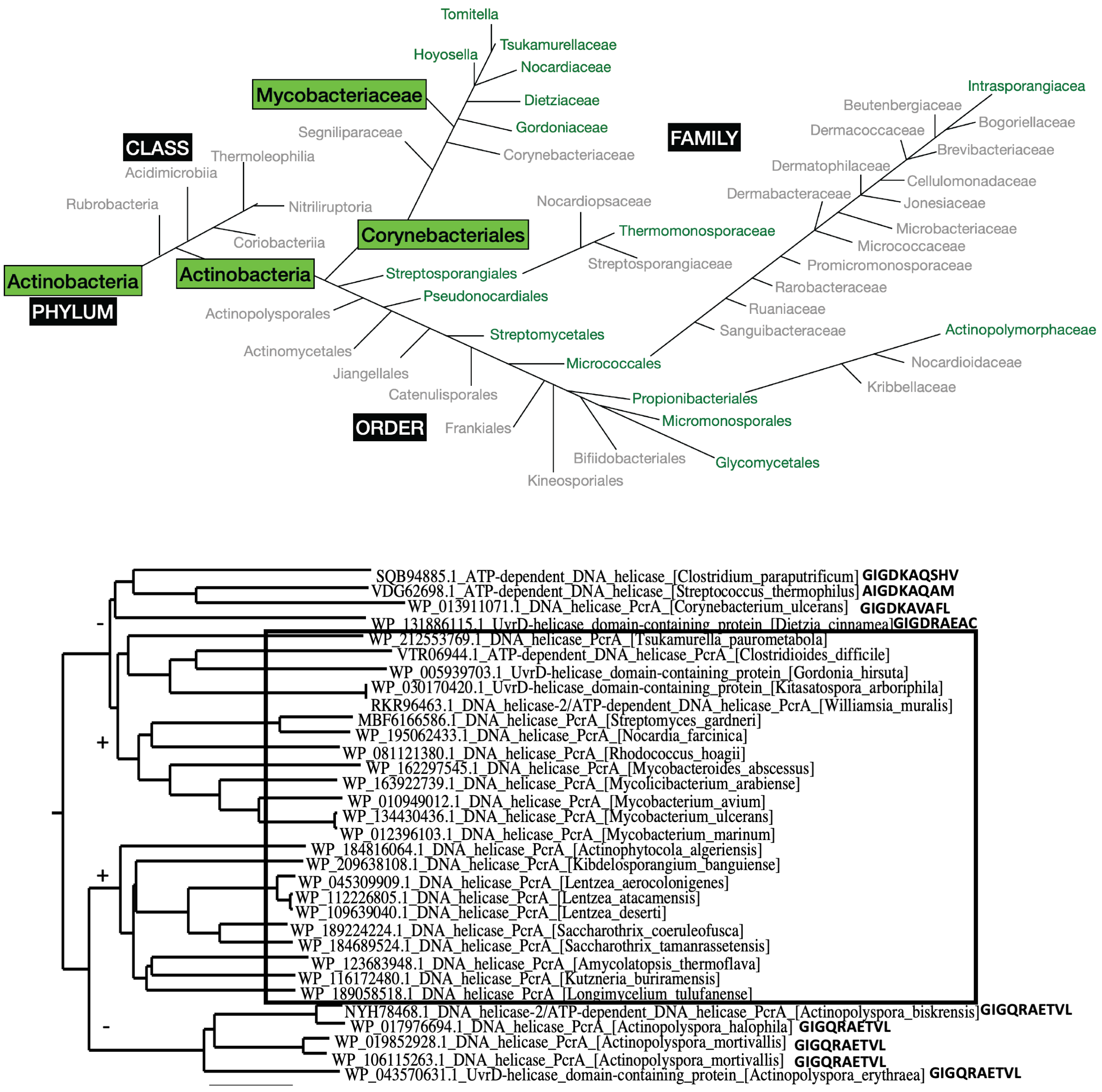


**Supplemental Figure 8: A more detailed look at the distribution of the 2B cysteine across the orders and families of the Actinobacteria class.** In the top panel, groups containing at least one organism with the 2B cysteine in the same position as UvrD1 are indicated in green. The boxed green labels refer to the family in which *Mtb* is found. Those that do not contain the 2B cysteine are indicated in gray. In the bottom panel a section of a larger phylogenic tree constructed using the UvrD1 sequence in different groups of actinobacteria and firmicutes. The boxed species all contain the 2B cysteine and the surrounding conserved residues as in Fig. 3B. The full tree can be seen on the next page.

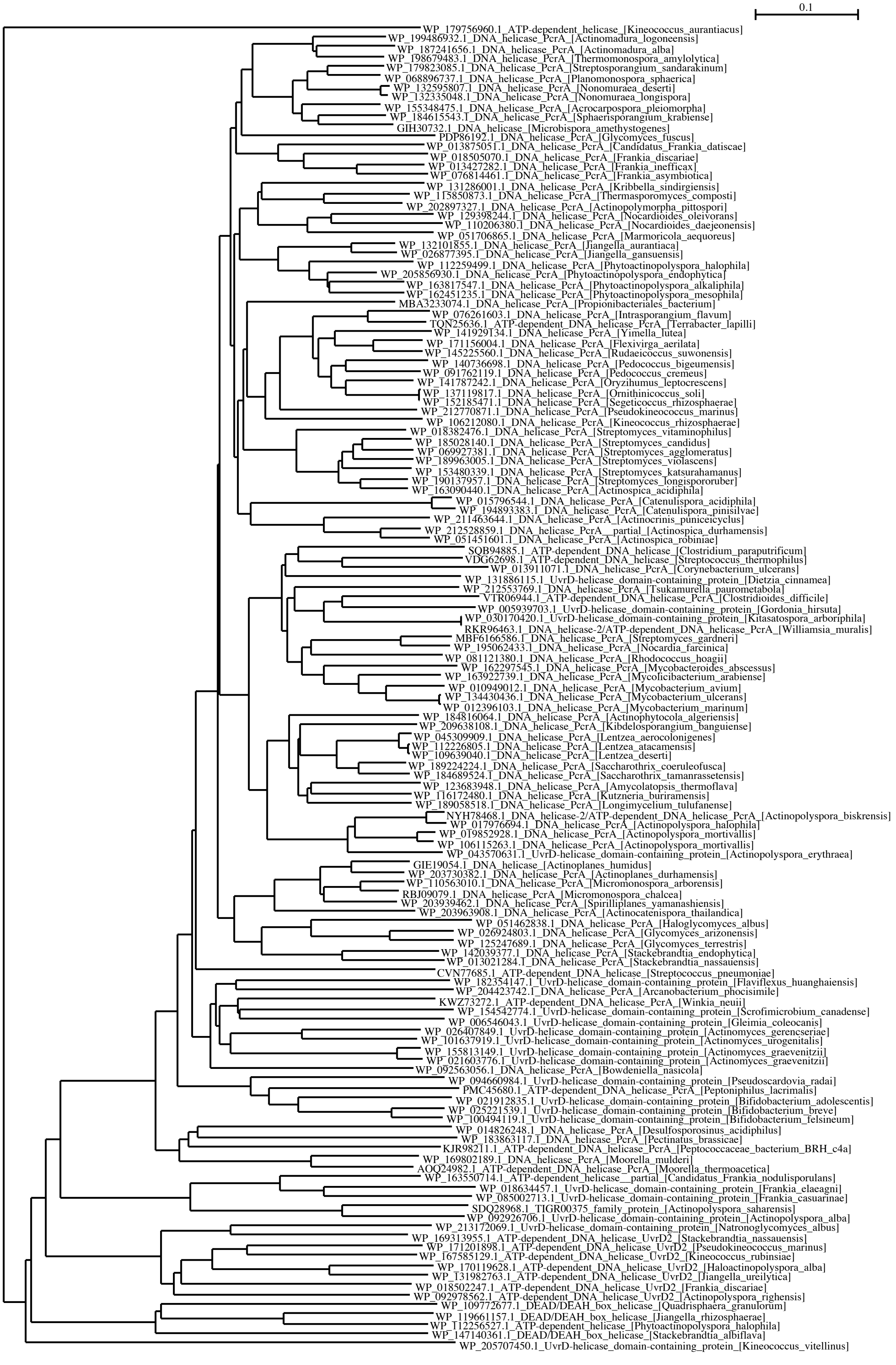
0.1
WP_179756960.1_ATP-dependent_helicase_[Kineococcus_aurantiacus]
WP_199486932.1_DNA_helicase_PcrA_[Actinomadura_logoneensis]
WP_187241656.1_DNA_helicase_PcrA_[Actinomadura_alba]
WP_198679483.1_DNA_helicase_PcrA_[Thermomonospora_amylolytica]
WP_179823085.1_DNA_helicase_PcrA_[Streptosporangium_sandarakinum]
WP_068896737.1_DNA_helicase_PcrA_[Planomonospora_sphaerica]
WP_132595807.1_DNA_helicase_PcrA_[Nonomuraea_deserti]
WP_132335048.1_DNA_helicase_PcrA_[Nonomuraea_longispora]
WP_155348475.1_DNA_helicase_PcrA_[Acrocarpospora_pleiomorpha]
WP_184615543.1_DNA_helicase_PcrA_[Sphaerisporangium_krabiense]
GIH30732.1_DNA_helicase_[Microbispora_amethystogenes]
PDP86192.1_DNA_helicase_PcrA_[Glycomyces_fuscus]
WP_013875051.1_DNA_helicase_PcrA_[Candidatus_Frankia_datiscae]
WP_018505070.1_DNA_helicase_PcrA_[Frankia_discariae]
WP_013427282.1_DNA_helicase_PcrA_[Frankia_inefficax]
WP_076814461.1_DNA_helicase_PcrA_[Frankia_asymbiotica]
WP_131286001.1_DNA_helicase_PcrA_[Kribbella_sindirgiensis]
WP_115850873.1_DNA_helicase_PcrA_[Thermasporomyces_composti]
WP_202897327.1_DNA_helicase_PcrA_[Actinopolymorpha_pittospori]
WP_129398244.1_DNA_helicase_PcrA_[Nocardioides_oleivorans]
WP_110206380.1_DNA_helicase_PcrA_[Nocardioides_daejeonensis]
WP_051706865.1_DNA_helicase_PcrA_[Marmoricola_aequoreus]
WP_132101855.1_DNA_helicase_PcrA_[Jiangella_aurantiaca]
WP_026877395.1_DNA_helicase_PcrA_[Jiangella_gansuensis]
WP_112259499.1_DNA_helicase_PcrA_[Phytoactinopolyspora_halophila]
WP_205856930.1_DNA_helicase_PcrA_[Phytoactinopolyspora_endophytica]
WP_163817547.1_DNA_helicase_PcrA_[Phytoactinopolyspora_alkaliphila]
WP_162451235.1_DNA_helicase_PcrA_[Phytoactinopolyspora_mesophila]
MBA3233074.1_DNA_helicase_PcrA_[Propionibacteriales_bacterium]
WP_076261603.1_DNA_helicase_PcrA_[Intrasporangium_flavum]
TQN25636.1_ATP-dependent_DNA_helicase_PcrA_[Terrabacter_lapilli]
WP_141929134.1_DNA_helicase_PcrA_[Yimella_lutea]
WP_171156004.1_DNA_helicase_PcrA_[Flexivirga_aerilata]
WP_145225560.1_DNA_helicase_PcrA_[Rudaeicoccus_suwonensis]
WP_140736698.1_DNA_helicase_PcrA_[Pedococcus_bigeumensis]
WP_091762119.1_DNA_helicase_PcrA_[Pedococcus_cremeus]
WP_141787242.1_DNA_helicase_PcrA_[Oryzihumus_leptocrescens]
WP_137119817.1_DNA_helicase_PcrA_[Ornithinicoccus_soli]
WP_152185471.1_DNA_helicase_PcrA_[Segeticoccus_rhizosphaerae]
WP_212770871.1_DNA_helicase_PcrA_[Pseudokineococcus_marinus]
WP_106212080.1_DNA_helicase_PcrA_[Kineococcus_rhizosphaerae]
WP_018382476.1_DNA_helicase_PcrA_[Streptomyces_vitaminophilus]
WP_185028140.1_DNA_helicase_PcrA_[Streptomyces_candidus]
WP_069927381.1_DNA_helicase_PcrA_[Streptomyces_agglomeratus]
WP_189963005.1_DNA_helicase_PcrA_[Streptomyces_violascens]
WP_153480339.1_DNA_helicase_PcrA_[Streptomyces_katsurahamanus]
WP_190137957.1_DNA_helicase_PcrA_[Streptomyces_longispororuber]
WP_163090440.1_DNA_helicase_PcrA_[Actinospica_acidiphila]
WP_015796544.1_DNA_helicase_PcrA_[Catenulispora_acidiphila]
WP_194893383.1_DNA_helicase_PcrA_[Catenulispora_pinisilvae]
WP_211463644.1_DNA_helicase_PcrA_[Actinocrinis_puniceicyclus]
WP_212528859.1_DNA_helicase_PcrA__partial_[Actinospica_durhamensis]
WP_051451601.1_DNA_helicase_PcrA_[Actinospica_robiniae]
SQB94885.1_ATP-dependent_DNA_helicase_[Clostridium_paraputrificum]
VDG62698.1_ATP-dependent_DNA_helicase_[Streptococcus_thermophilus]
WP_013911071.1_DNA_helicase_PcrA_[Corynebacterium_ulcerans]
WP_131886115.1_UvrD-helicase_domain-containing_protein_[Dietzia_cinnamea]
WP_212553769.1_DNA_helicase_PcrA_[Tsukamurella_paurometabola]
VTR06944.1_ATP-dependent_DNA_helicase_PcrA_[Clostridioides_difficile]
WP_005939703.1_UvrD-helicase_domain-containing_protein_[Gordonia_hirsuta]
WP_030170420.1_UvrD-helicase_domain-containing_protein_[Kitasatospora_arboriphila]
RKR96463.1_DNA_helicase-2/ATP-dependent_DNA_helicase_PcrA_[Williamsia_muralis]
MBF6166586.1_DNA_helicase_PcrA_[Streptomyces_gardneri]
WP_195062433.1_DNA_helicase_PcrA_[Nocardia_farcinica]
WP_081121380.1_DNA_helicase_PcrA_[Rhodococcus_hoagii]
WP_162297545.1_DNA_helicase_PcrA_[Mycobacteroides_abscessus]
WP_163922739.1_DNA_helicase_PcrA_[Mycolicibacterium_arabiense]
WP_010949012.1_DNA_helicase_PcrA_[Mycobacterium_avium]
WP_134430436.1_DNA_helicase_PcrA_[Mycobacterium_ulcerans]
WP_012396103.1_DNA_helicase_PcrA_[Mycobacterium_marinum]
WP_184816064.1_DNA_helicase_PcrA_[Actinophytocola_algeriensis]
WP_209638108.1_DNA_helicase_PcrA_[Kibdelosporangium_banguiense]
WP_045309909.1_DNA_helicase_PcrA_[Lentzea_aerocolonigenes]
WP_112226805.1_DNA_helicase_PcrA_[Lentzea_atacamensis]
WP_109639040.1_DNA_helicase_PcrA_[Lentzea_deserti]
WP_189224224.1_DNA_helicase_PcrA_[Saccharothrix_coeruleofusca]
WP_184689524.1_DNA_helicase_PcrA_[Saccharothrix_tamanrassetensis]
WP_123683948.1_DNA_helicase_PcrA_[Amycolatopsis_thermoflava]
WP_116172480.1_DNA_helicase_PcrA_[Kutzneria_buriramensis]
WP_189058518.1_DNA_helicase_PcrA_[Longimycelium_tulufanense]
NYH78468.1_DNA_helicase-2/ATP-dependent_DNA_helicase_PcrA_[Actinopolyspora_biskrensis]
WP_017976694.1_DNA_helicase_PcrA_[Actinopolyspora_halophila]
WP_019852928.1_DNA_helicase_PcrA_[Actinopolyspora_mortivallis]
WP_106115263.1_DNA_helicase_PcrA_[Actinopolyspora_mortivallis]
WP_043570631.1_UvrD-helicase_domain-containing_protein_[Actinopolyspora_erythraea]
GIE19054.1_DNA_helicase_[Actinoplanes_humidus]
WP_203730382.1_DNA_helicase_PcrA_[Actinoplanes_durhamensis]
WP_110563010.1_DNA_helicase_PcrA_[Micromonospora_arborensis]
RBJ09079.1_DNA_helicase_PcrA_[Micromonospora_chalcea]
WP_203939462.1_DNA_helicase_PcrA_[Spirilliplanes_yamanashiensis]
WP_203963908.1_DNA_helicase_PcrA_[Actinocatenispora_thailandica]
WP_051462838.1_DNA_helicase_PcrA_[Haloglycomyces_albus]
WP_026924803.1_DNA_helicase_PcrA_[Glycomyces_arizonensis]
WP_125247689.1_DNA_helicase_PcrA_[Glycomyces_terrestris]
WP_142039377.1_DNA_helicase_PcrA_[Stackebrandtia_endophytica]
WP_013021284.1_DNA_helicase_PcrA_[Stackebrandtia_nassauensis]
CVN77685.1_ATP-dependent_DNA_helicase_[Streptococcus_pneumoniae]
WP_182354147.1_UvrD-helicase_domain-containing_protein_[Flaviflexus_huanghaiensis]
WP_204423742.1_DNA_helicase_PcrA_[Arcanobacterium_phocisimile]
KWZ73272.1_ATP-dependent_DNA_helicase_PcrA_[Winkia_neuii]
WP_154542774.1_UvrD-helicase_domain-containing_protein_[Scrofimicrobium_canadense]
WP_006546043.1_UvrD-helicase_domain-containing_protein_[Gleimia_coleocanis]
WP_026407849.1_UvrD-helicase_domain-containing_protein_[Actinomyces_gerencseriae]
WP_101637919.1_UvrD-helicase_domain-containing_protein_[Actinomyces_urogenitalis]
WP_155813149.1_UvrD-helicase_domain-containing_protein_[Actinomyces_graevenitzii]
WP_021603776.1_UvrD-helicase_domain-containing_protein_[Actinomyces_graevenitzii]
WP_092563056.1_DNA_helicase_PcrA_[Bowdeniella_nasicola]
WP_094660984.1_UvrD-helicase_domain-containing_protein_[Pseudoscardovia_radai]
PMC45680.1_ATP-dependent_DNA_helicase_PcrA_[Peptoniphilus_lacrimalis]
WP_021912835.1_UvrD-helicase_domain-containing_protein_[Bifidobacterium_adolescentis]
WP_025221539.1_UvrD-helicase_domain-containing_protein_[Bifidobacterium_breve]
WP_100494119.1_UvrD-helicase_domain-containing_protein_[Bifidobacterium_felsineum]
WP_014826248.1_DNA_helicase_PcrA_[Desulfosporosinus_acidiphilus]
WP_183863117.1_DNA_helicase_PcrA_[Pectinatus_brassicae]
KJR98211.1_ATP-dependent_DNA_helicase_PcrA_[Peptococcaceae_bacterium_BRH_c4a]
WP_169802189.1_DNA_helicase_PcrA_[Moorella_mulderi]
AOQ24982.1_ATP-dependent_DNA_helicase_PcrA_[Moorella_thermoacetica]
WP_163550714.1_ATP-dependent_helicase__partial_[Candidatus_Frankia_nodulisporulans]
WP_018634457.1_UvrD-helicase_domain-containing_protein_[Frankia_elaeagni]
WP_085002713.1_UvrD-helicase_domain-containing_protein_[Frankia_casuarinae]
SDQ28968.1_TIGR00375_family_protein_[Actinopolyspora_saharensis]
WP_092926706.1_UvrD-helicase_domain-containing_protein_[Actinopolyspora_alba]
WP_213172069.1_UvrD-helicase_domain-containing_protein_[Natronoglycomyces_albus]
WP_169313955.1_ATP-dependent_DNA_helicase_UvrD2_[Stackebrandtia_nassauensis]
WP_171201898.1_ATP-dependent_DNA_helicase_UvrD2_[Pseudokineococcus_marinus]
WP_167585129.1_ATP-dependent_DNA_helicase_UvrD2_[Kineococcus_rubinsiae]
WP_170119628.1_ATP-dependent_DNA_helicase_UvrD2_[Haloactinopolyspora_alba]
WP_131982763.1_ATP-dependent_DNA_helicase_UvrD2_[Jiangella_ureilytica]
WP_018502247.1_ATP-dependent_DNA_helicase_UvrD2_[Frankia_discariae]
WP_092978562.1_ATP-dependent_DNA_helicase_UvrD2_[Actinopolyspora_righensis]
WP_109772677.1_DEAD/DEAH_box_helicase_[Quadrisphaera_granulorum]
WP_119661157.1_DEAD/DEAH_box_helicase_[Jiangella_rhizosphaerae]
WP_112256527.1_ATP-dependent_helicase_[Phytoactinopolyspora_halophila]
WP_147140361.1_DEAD/DEAH_box_helicase_[Stackebrandtia_albiflava]
WP_205707450.1_UvrD-helicase_domain-containing_protein_[Kineococcus_vitellinus]

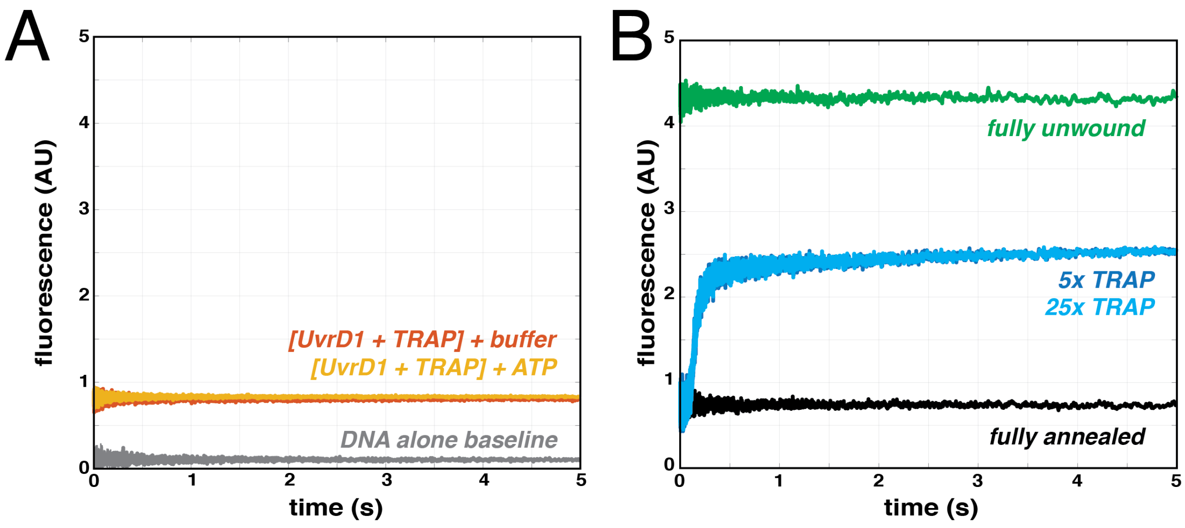


**Supplemental Figure 9: DNA unwinding controls and calculation of fraction unwound. (A)** When TRAP DNA is equilibrated with UvrD1 and the DNA unwinding template and shot against ATP (yellow), the signal is constant and identical to that in the absence of ATP (orange). Note that the addition of UvrD1 does lead to a slight increase in the fluorescence signal compared to DNA alone (gray). **(B)** The fluorescence signals for traces in the presence of the same UvrD1 concentration including the fully unwound template (green), the fully annealed template (black), and two experimental traces with different molar ratios of TRAP (5X (blue) and 25X (cyan) relative to 200 nM protein). The fraction unwound was calculated as (signal – negative control) / (positive control – negative control) as described in the Methods.

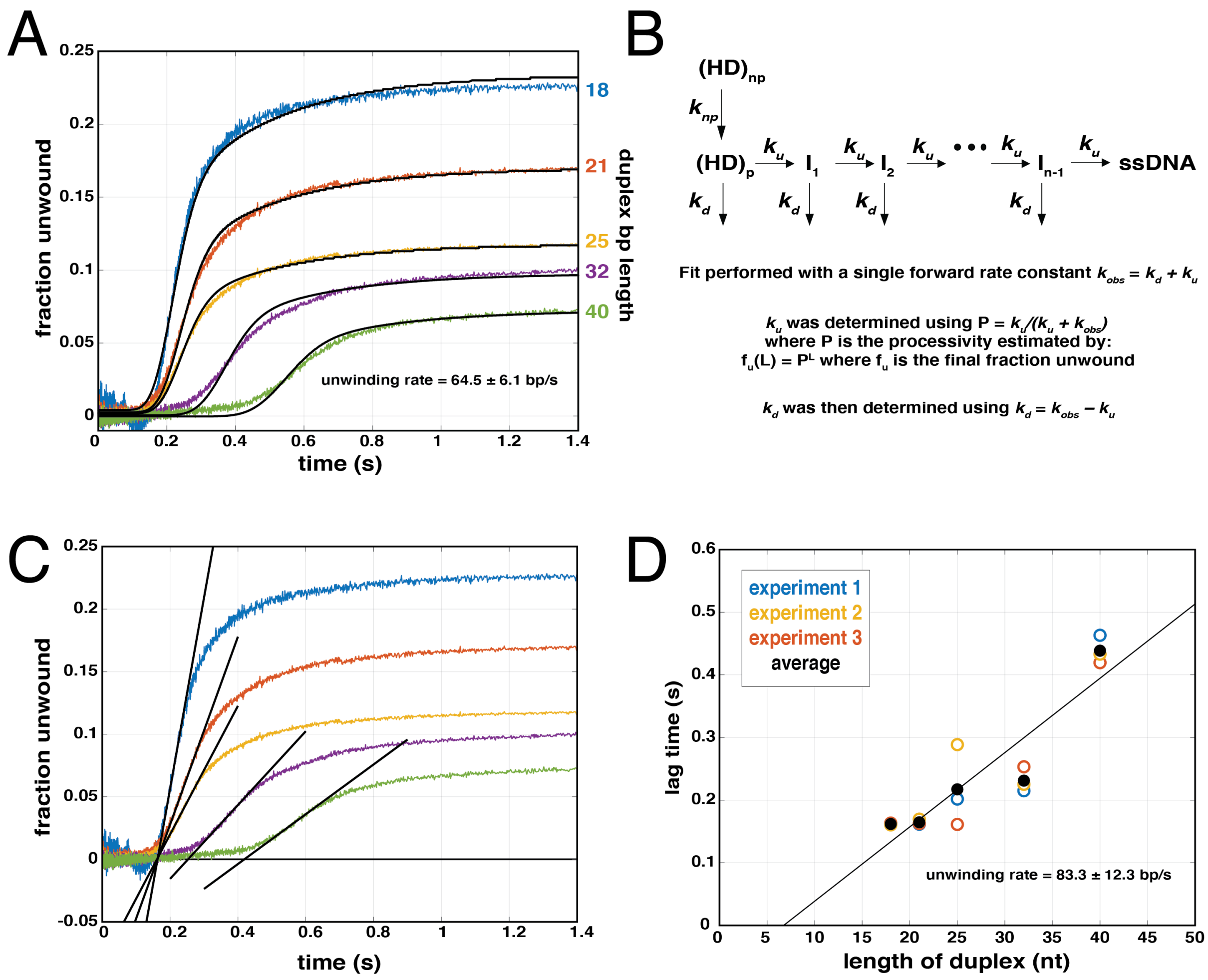


**Supplemental Figure 10: DNA unwinding as a function of duplex length.** DNA templates of different duplex base-pair lengths (18, 21, 25, 32, 40) were used in the unwinding assay under identical conditions to those in the main text (Fig. 4A) for *Mtb* UvrD1 in the absence of DTT. **(A)** As expected for a processive unwinding mechanism, the kinetics and the final fraction unwound in a single turnover reaction decreased with increasing length of DNA duplex. Examples fits of to an n-step kinetic model including a non-productive fraction are shown (black) and lead to an estimate of an unwinding rate of 64.5 ± 6.1 bp/s, a dissociation rate of 3.2 ± 0.9 $s^{-1}$, and a non-productive isomerization rate of 3.5 ± 0.2 $s^{-1}$. The fit parameters for three separate experiments can be seen in **Supplemental Table 2**. **(B)** The kinetic scheme used to fit the traces in (A) is shown. A productive complex $(HD)_p$ can take an unwinding step ($k_u$) or dissociate ($k_d$) at each position until arriving at a completely unwound template (ssDNA). A non-productive complex $(HD)_{np}$ must first isomerize to a productive complex ($k_{np}$) before beginning to unwind. **(C)** The same data as in (A) showing the estimation of lag times. Lag times were taken as the intercept of the linear portions of the traces with the X-axis as shown. **(D)** The lag times were measured for each length in each of the three experiments (blue, yellow, and orange circles). The average of these lag times (black circles) was used to fit a line who's inverse gives an estimate of the unwinding rate of 83.3 ± 12.3 bp/s.

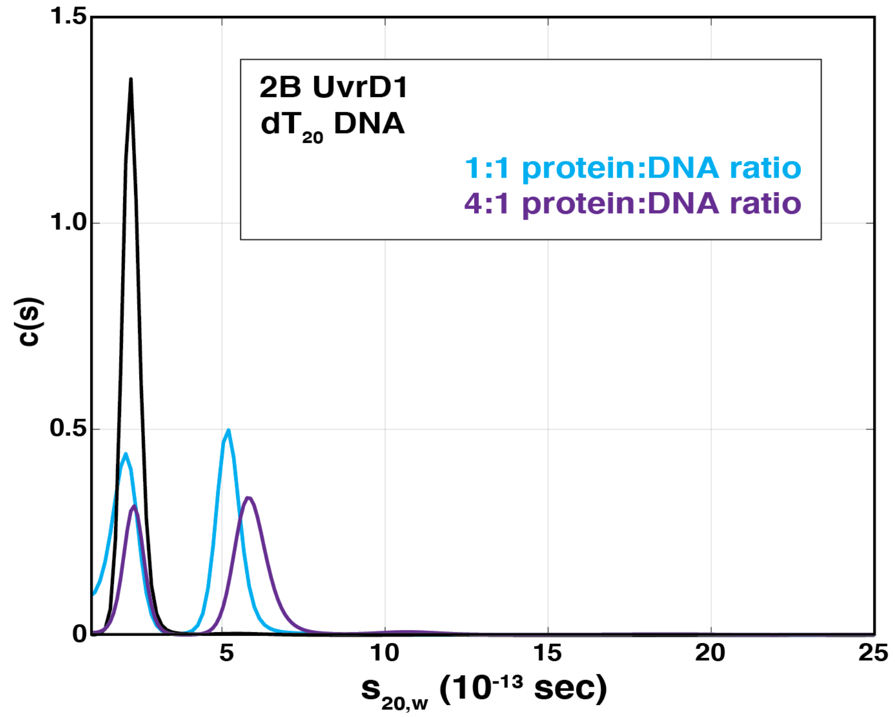


**Supplemental Figure 11: Excess monomer does not result in two peaks in sedimentation velocity.** Sedimentation velocity c(s) distributions obtained by monitoring Cy5 fluorescence in Buffer A at 75 mM NaCl in the absence of DTT for $dT_{20}$ DNA alone (black) and the 2B mutant either at a 4X molar excess of protein (purple) or where protein and DNA are at a 1:1 molar stoichiometry (cyan). In both cases a single protein-DNA peak was observed and no sedimenting species near an $s_{20,w}$ value of 8, as measured for the WT dimer DNA bound complex (Fig. 5B, Supp. Table 3), was detected for either protein:DNA ratio. The shift in the protein-bound peak here may be due to rapid fluctuations between a single monomer and two monomers bound to the tail without formation of the stable, active dimer. Alternatively, at these higher protein:DNA ratios, the active dimer may form, but would be too short lived to result in unwiding.

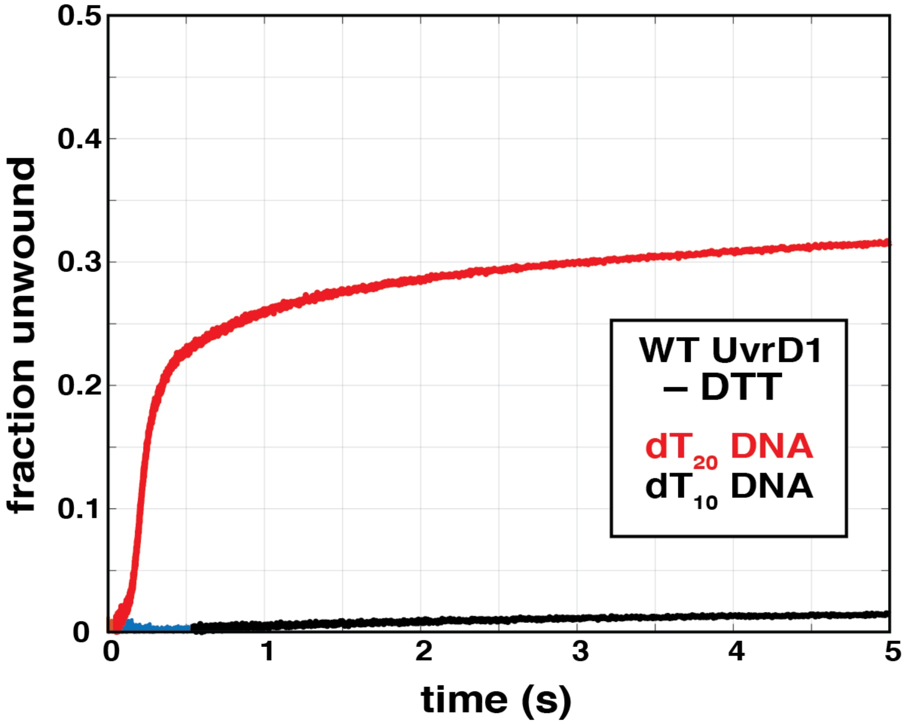


**Supplemental Figure 12: Comparison of fraction DNA unwound with different lengths of ssDNA tails as a function of time.** Experiments here are performed under identical conditions to those in Fig. 4B for WT UvrD1 in the absence of DTT. In contrast to the case with a $dT_{20}$ tail (red) that permits both the monomer and dimer species to bind the unwinding template (Fig. 5B), minimal DNA unwinding is observed with a $dT_{10}$ tail (black), where only the monomer is observed bound to DNA (Fig. 5B). These results indicate that both monomers of the UvrD1 dimer must interact with the DNA to form an unwinding-competent species.

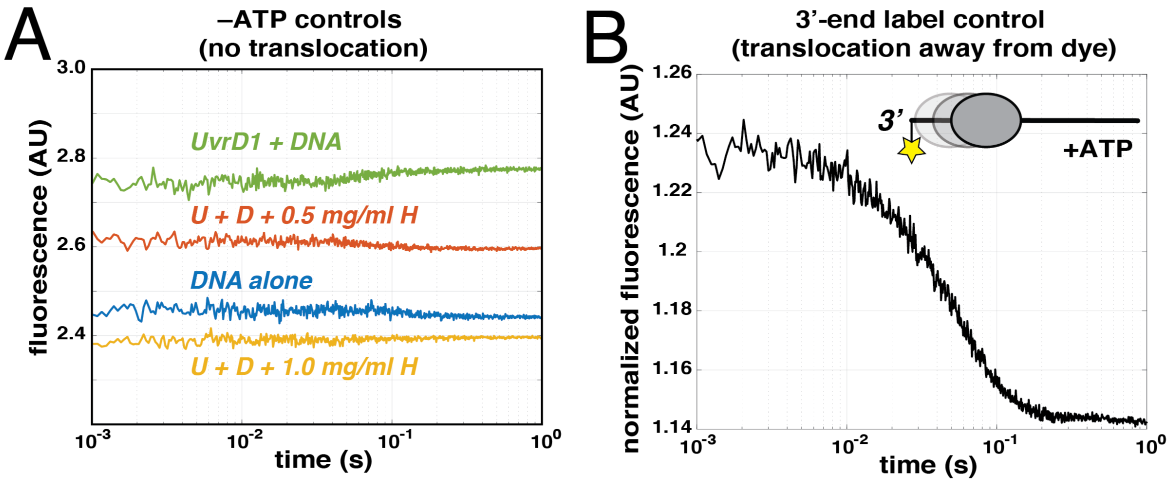


**Supplemental Figure 13: Heparin controls for establishing single round conditions in translocation assays.** Experiments were conducted in Buffer A with 75 mM NaCl at 25 °C in the absence of DTT. **(A)** Preincubation of UvrD1 and DNA (green) in the absence of ATP and $Mg^{2+}$ leads to a small increase in fluorescence relative to DNA alone (blue). Addition of 0.5 mg/ml heparin during the preincubation lowers this signal and 1.0 mg/ml heparin (yellow) leads to a fluorescence similar to DNA alone suggesting it completely prevents UvrD1-DNA binding at this concentration. **(B)** Using a 3'-end labeled DNA template instead of a 5'-end labeled template results in a decrease in fluorescence upon mixing pre-bound UvrD1 with ATP and $Mg^{2+}$. Fluorescence has been normalized by dividing by the signal from a DNA only trace.

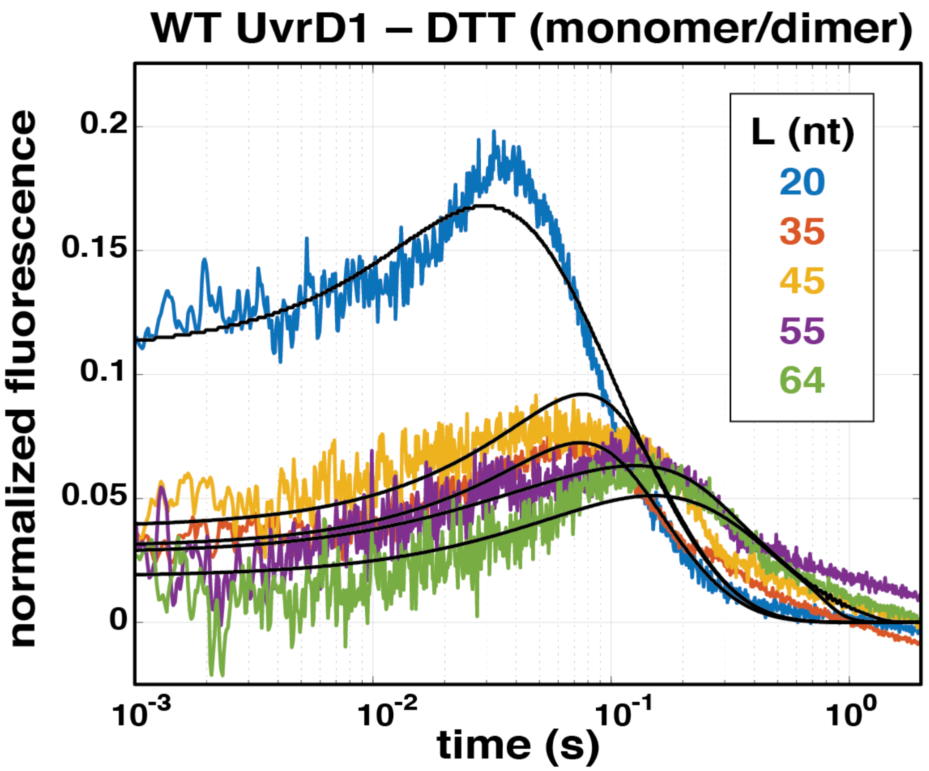


**Supplementary Figure S14: Translocation traces from UvrD1 in the absence of reducing agent.** The presence of both monomeric and dimeric species results in translocation kinetics distinct from that of monomer alone (see main text Fig. 6). Fit parameters to an n-step sequential model with two populations of translocating species can be found in Supplemental Table 4. These traces were normalized to the average of last ten plateau values observed in the raw data at each DNA length, for ease in fitting and visualization.

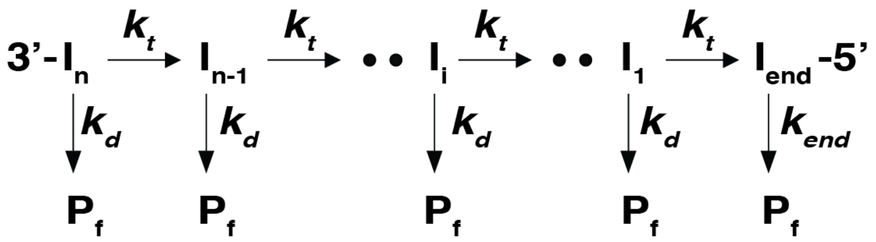


**Supplemental Figure 15: N-step kinetic scheme for fitting translocation data.** Translocation steps forward occurs with a rate $k_t$ while dissociation from internal sites occurs with a rate $k_d$ and dissociation from the 5'-end of the template occurs with a rate $k_{end}$.

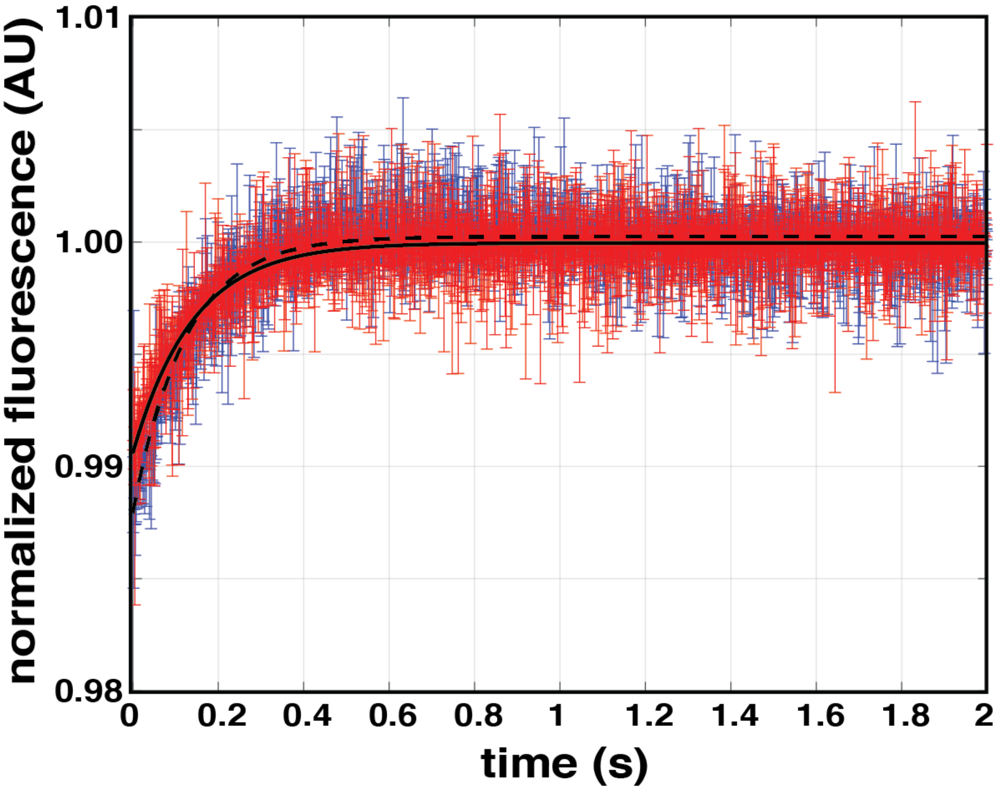


**Supplemental Figure 16: Dissociation kinetics of WT UvrD1 in the presence and absence of DTT measured with tryptophan fluorescence.** UvrD1 dissociation kinetics was measured from $(dT)_{100}$ for WT+DTT (blue) and poly$(dT)_{964}$ for WT-DTT (red) (Methods). Excess UvrD1 was prebound to the dT DNA oligos and mixed with identical concentrations of ATP, heparin, and $Mg^{2+}$ as those in the translocation assays. Mixing led to UvrD1 dissociation as monitored by an increase in tryptophan fluorescence measured using 290nm excitation and 305nm long pass emission filter. Fitting both traces with a single exponential function (black lines) yielded similar estimates for the observed dissociation rates for both the +DTT condition ($8.1 \pm 0.2$ $s^{-1}$) and the -DTT condition ($7.1 \pm 0.2$ $s^{-1}$). These values were used to constrain the observed dissociation rates for the monomer and dimer, respectively for the global fitting of translocation traces.

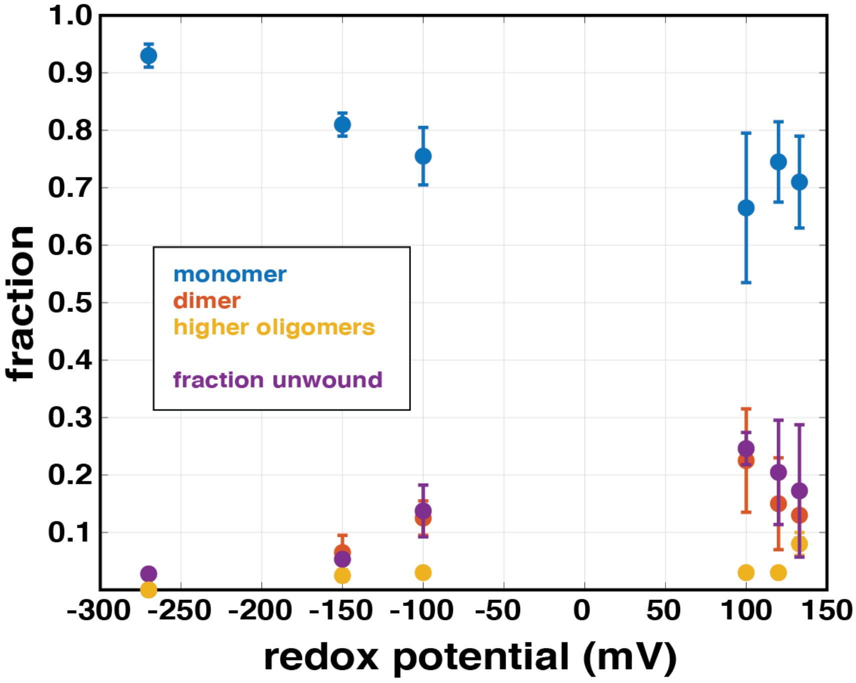


**Supplemental Figure 17: Dimer fraction correlates with fraction of DNA unwound.** The equivalent of Fig. 7 in the main text but using 2 μM UvrD1 1A1B double mutant.

**Full Genus names for species listed in Fig. 3B**

*Escherichia coli*
*Escherichia coli*
*Bacillus subtilis*
*Mycobacterium tuberculosis*
*Clostridioides difficile*
*Gordonia bronchialis*
*Dietzia cinname*
*Tsukamurella paurometabola*
*Hoyosella rhizosphaerae*
*Tomotella caverna*
*Mycobacterium ulcerans*
*Rhodococccus hoagie*
*Streptomyces regensis*
*Stackebrandtia nassauensis*
*Actinocatenispora thailandica*
*Longimycelium tulufanens*
*Actinopolymorpha pittospori*
*Terrabacter lapilli*
*Actinomadura logoneensis*

| condition | MW (Da) calculated | MW (Da) measured | $s_{20,w}$ measured | $f/f_0$ measured | fraction oligomer |
|---|---|---|---|---|---|
| **Monomers** | | | | | |
| WT –DTT | 85,049 | 77,000 | 4.72 | 1.17 | 0.3 |
| WT +DTT | 85,049 | 82,600 | 4.72 | 1.17 | 0.9 |
| WT 400 mM NaCl –DTT | 85,049 | 68,100 | 4.22 | 1.2 | 0.6 |
| WT 400 mM NaCl +DTT | 85,049 | 74,600 | 4.34 | 1.19 | 0.92 |
| 2B –DTT | 85,017 | 88,100 | 4.61 | 1.17 | 0.9 |
| 2B +DTT | 85,017 | 83,000 | 4.51 | 1.19 | 0.92 |
| 1A1B +DTT | 85,146 | 86,200 | 4.72 | 1.2 | 0.94 |
| 1A1B –DTT | 85,146 | 81,000 | 5.14 | 1.21 | 0.3 |
| **Dimers** | | | | | |
| WT –DTT | 170,098 | 180,020 | 7.55 | 1.17 | 0.3 |
| 1A1B –DTT | 170,292 | 185,000 | 7.55 | 1.17 | 0.6 |
| WT 400mM NaCl –DTT | 170,098 | 131,000 | 6.79 | 1.2 | 0.3 |
| **Higher-order Oligomers** | | | | | |
| WT –DTT | 340,196 | 367,000 | 9.54 | 1.17 | 0.33 |
| 1A1B +DTT | 340,584 | 373,000 | 14.5 | 1.2 | 0.015 |
| 1A1B –DTT | 340,584 | 333,000 | 14.5 | 1.17 | 0.05 |
| WT 400 mM NaCl –DTT | 340,196 | 298,000 | 14.3 | 1.2 | 0.03 |
| 2B –DTT | 340,068 | 379,000 | 13.63 | 1.17 | 0.03 |
| 2B +DTT | 340,068 | 380,000 | 13.62 | 1.19 | 0.028 |

**Supplemental Table 1: Summary of sedimentation velocity results obtained in the absence of DNA.** The hydrodynamic properties of each oligomeric form for both wild-type and cysteine mutant constructs of *Mtb* UvrD1 at 2.5 µM protein concentration. Unless specified, all quantifications presented were obtained in Buffer A with 75 mM NaCl at 25 $^{o}$C.

| Parameter replicate: | first | second | third | average |
|---|---|---|---|---|
| $k_u$ ($s^{-1}$) | 58.5 ± 1.4 | 76.0 ± 1.1 | 82.6 ± 1.3 | 72.4 ± 12.5 |
| m (bp/step) | 1.01 ± 0.06 | 0.83 ± 0.18 | 0.84 ± 0.38 | 0.89 ± 0.10 |
| $mk_u$ (bp-$s^{-1}$) | 59.1 ± 3.8 | 63.1 ± 13.7 | 69.4 ± 28.6 | 63.9 ± 5.2 |
| $k_d$ ($s^{-1}$) | 2.3 ± 1.6 | 4.0 ± 1.4 | 3.4 ± 1.9 | 3.2 ± 0.9 |
| $k_{np}$ ($s^{-1}$) | 3.70 ± 0.02 | 3.30 ± 0.03 | 3.60 ± 0.05 | 3.53 ± 0.21 |
| $x_p$ | 0.58 ± 0.002 | 0.66 ± 0.003 | 0.56 ± 0.004 | 0.60 ± 0.05 |
| Processivity (P) | 0.95 ± 0.02 | 0.95 ± 0.007 | 0.95 ± 0.002 | 0.95 ± 0.01 |
| bp unwound per binding event | 20 ± 10 | 20 ± 3 | 20 ± 1 | 20.0 ± 0.01 |
| $Chi^2$ | 0.15 | 0.27 | 0.23 | |

**Supplemental Table 2: Parameter estimates from fits to unwinding data.** Unwinding parameters obtained from fits of data shown in Fig. S10A to the model in Fig. S10B. Unwinding rate ($k_u$), kinetic step-size (m), overall unwinding rate ($mk_u$), dissociation rate ($k_d$), non-productive isomerization rate ($k_{np}$), fraction productive complexes ($x_p$), processivity (p) and base-pairs unwound per binding event are shown as independently obtained from three replicate experiments along with estimated fit errors. The average of each parameter is shown in the last column along with the standard deviation of the three measurements.

| Species | MW (Da) calculated | MW (Da) measured | $s_{20,w}$ measured | $f/f_0$ measured | fraction DNA/ oligomer | v (ml/g) calculated |
|---|---|---|---|---|---|---|
| 3'-dT(10)-ds18-cy5 | 14500 | 10600 | 2.41 | 1.21 | 0.97 | 0.563 |
| 3'-dT(20)-ds18-cy5 | 19000 | 12800 | 2.2 | 1.47 | 0.92 | 0.563 |
| $(2B\text{-}UvrD1)_1$ + 3'-dT(20)-ds18-cy5 | 104008 | 90800 | 5.2 | 1.19 | 0.47 | 0.708 |
| $(WT\text{-}UvrD1)_1$ + 3'-dT(10)-ds18-cy5 | 99508 | 104000 | 5.55 | 1.21 | 0.35 | 0.709 |
| $(WT\text{-}UvrD1)_1$ + 3'-dT(20)-ds18-cy5 | 104008 | 96300 | 5.55 | 1.18 | 0.31 | 0.725 |
| $(WT\text{-}UvrD1)_2$ + 3'-dT(20)-ds18-cy5 | 199000 | 162000 | 8.07 | 1.18 | 0.31 | 0.725 |

**Supplemental Table 3: Summary of sedimentation velocity results obtained in the presence of DNA.** The hydrodynamic properties for labeled DNA alone and DNA complexes with either monomer or dimer WT and 2B mutant UvrD1 at 1.5 or 4 µM protein concentration in case of 2B mutant. All quantifications presented were obtained in Buffer A with 75 mM NaCl at 25 $^{o}$C in the absence of DTT.

| parameter | WT +DTT (average) | 2B –DTT (average) | WT –DTT (average) |
|---|---|---|---|
| $k_t$ ($s^{-1}$) | 43 ± 6 | 43 ± 1 | 40 |
| $k_d$ ($s^{-1}$) | 3.8 ± 0.3 | 4.1 ± 0.1 | 4.1 |
| $k_{end}$ ($s^{-1}$) | 63 ± 3 | 61 ± 4 | 61 |
| r | 2.4 ± 0.5 | 2.6 ± 0.2 | 2.6 |
| m (nt) | 3.0 ± 0.5 | 3.0 ± 0.7 | 0.8 ± 0.1 |
| $d_{app}$ (nt) | 13 ± 1 | 12 ± 1 | 23 ± 5 |
| $mk_t$ (nt/s) | 120 ± 5 | 130 ± 10 | 71 ± 15 |
| $f_m$ | - | - | 0.3 |
| $f_d$ | - | - | 0.7 |
| $k_{t2}$ ($s^{-1}$) | - | - | 88 ± 6 |
| $k_{d2,obs}$ ($s^{-1}$) | - | - | 2 ± 1 |
| $k_{end2,obs}$ ($s^{-1}$) | - | - | 7 ± 3 |
| r2 | - | - | 0.5 ± 0.1 |
| processivity | 32 ± 1 | 32 ± 5 | 53 ± 43 |
| $Chi^2$ | 0.4 ± 0.1 | 0.2 ± 0.1 | 1 ± 0.3 |

**Supplemental Table 4: Parameter estimates from fits of translocation data.** Parameters are as follows: $k_t$ is translocation rate, $k_d$ is dissociation rate from internal sites, $k_{end}$ is the dissociation rate from the ends, r is the binding probability ratio (random/end), m is kinetic step size, $d_{app}$ is the apparent contact size, $mk_t$ is the macroscopic translocation rate, $f_m$ is the fraction monomer and $f_d$ is the fraction dimer population for the mixed fits where both monomer and dimer populations are present, $k_{t2}$ is the translocation rate for the dimer, $k_{d2}$ is the dissociation rate for the dimer, $k_{end2}$ is the rate of dissociation from the ends, r2 is the probability ratio (random/end) for the dimer, $Chi^2$ is the goodness of fit, and processivity is calculated from the rates of translocation and dissociation and is the average number of steps along the template taken prior to dissociation.

| Name | Sequence | Construct used for |
|---|---|---|
| BamH1 forward WT UvrD1 | 5'CCCGGATCCATGAGTGTGCACGCGACCG 3' | Wild type UvrD1 |
| Hind III reverse WT UvrD1 | 5'CGCAAGCTTTCAGAGCTTGGTGACAGGGGCGTGGTTG 3' | |
| 451 cysteine to alanine forward | 5'GATCGTGCCGAGGCGGCAGTGGCGGTGTACGCCGAGAAC 3' | 2B mutant UvrD1 (C451A) |
| 451 cysteine to alanine reverse | 5'GTTCTCGGCGTACACCGCCACTGCCGCCTCGGCACGATC 3' | |
| 269 cysteine to threonine forward | 5'CCCGGCGAGTTGACCGTCGTCGGGGATGCCG 3' | 1A1B double mutant UvrD1 (C107T/C269T) |
| 269 cysteine to threonine reverse | 5'CGGCATCCCCGACGACGGTCAACTCGCCGGG 3' | |
| 107 cysteine to threonine forward | 5'GACGTTTCACTCCACCACGGTGCGTTATCCTTGCGCAACC 3' | |
| 107 cysteine to threonine reverse | 5'GGTTGCGCAAGGATAACGCACCGTGGTGGAGTGAAACGTC 3' | |

**Supplementary Table 5: DNA primers used for creating mutant UvrD1 plasmids.**

| Length (nt) | Sequence | Sequences used for |
|---|---|---|
| 18 | 5'-GTT GGT CGG CAG CAG GGC-3' | Trap |
| 18 | 5'-GCC CTG CTG CCG ACC AAC -3' | Anisotropy assays |
| 38 | 5'-56-FAM GTT GGT CGG CAG CAG GGC-dT(20)3' | |
| 18 | 5'-GCC CTG CTG CCG ACC AAC-BHQ_2 -3' | Double-stranded DNA unwinding assays |
| 38 | 5'-cy5 GTT GGT CGG CAG CAG GGC-dT(20)3' | |
| 21 | 5′-GCC CTG CTG CCG ACC AAC GAT- (BHQ_2)-3′ | |
| 41 | 5'-cy5 ATC GTT GGT CGG CAG CAG GGC-dT(20)3' | |
| 25 | 5′-GCC CTG CTG CCG ACC AAC GAT GGT T- (BHQ_2)-3′ | |
| 45 | 5'-cy5 AA CC ATC GTT GGT CGG CAG CAG GGC-dT(20)3' | |
| 32 | 5′-GCC CTG CTG CCG ACC AAC GAT GGT TAC ATT CC (BHQ_2)-3′ | |
| 52 | 5'-cy5 GG AAT GTA ACC ATC GTT GGT CGG CAG CAG GGC-dT(20)3' | |
| 40 | 5′-GCC CTG CTG CCG ACC AAC GAT GGT TAC ATT CCC GCT GCT G (BHQ_2)-3′ | |
| 60 | 5'-cy5 C AGC AGC GGG AAT GTA ACC ATC GTT GGT CGG CAG CAG GGC-dT(20)3' | |
| 20 | 5'-cy3 dT(20)-3' | Single-stranded DNA translocation assays |
| 35 | 5'-cy3 dT(35)-3' | |
| 45 | 5'-cy3 dT(45)-3' | |
| 75 | 5'-cy3 dT(75)-3' | |
| 104 | 5'-cy3 dT(104)-3' | |

**Supplemental Table 6: Sequences of single-stranded DNA oligomers used for translocation and DNA unwinding studies.**
